# RNA Binding Protein Khdrbs1 Regulates Hematopoietic Stem and Progenitor Cell Emergence via Splicing

**DOI:** 10.1101/2024.11.01.621494

**Authors:** Ilana Karp, Teresa V. Bowman

## Abstract

Definitive hematopoietic stem and progenitor cells (HSPCs) arise through an endothelial-to-hematopoietic transition (EHT), whereby endothelial cells (ECs) acquire a hemogenic fate (HE) and convert to HSPCs. EHT regulators remain incompletely understood. Utilizing published RNA-seq data of ECs, HE/HSPCs, and HSPCs, we defined the alternative splicing landscape of zebrafish EHT and potential RNA binding proteins governing the process. Employing an F0 CRISPR-Cas9 mediated mutagenesis screen, we identified zebrafish homologs of KHDRBS1 as critical for embryonic HSPC formation. Mutagenesis of *khdrbs1a/b* led to diminished HSPCs and a profound lineage shift in differentiated cells, with the greatest decrease observed in *rag+* T-cells. Treatment of mutagenized embryos with the splicing modulator E7107 partially restored HSPCs to near wildtype levels indicating Khdrbs1 splicing regulation drives EHT. Consistently, Khdrbs1 binding motifs were the only ones enriched in EHT alternative splicing events. Identification of EHT-specific splicing regulation provides a deeper understanding of cell fate tuning by splicing.

## INTRODUCTION

Hematopoietic stem and progenitor cells (HSPCs) sustain lifelong hematopoiesis and are characterized by their ability to self-renew and differentiate into all blood cell lineages. The *de novo* production of HSPCs occurs during early embryonic development during a finite critical window through a process termed the endothelial-to-hematopoietic transition (EHT)^1–3^. The EHT consists of two distinct phases: 1) adaptation of a hemogenic transcription profile by a small population of endothelial cells (ECs) to form hemogenic endothelium (HE)^4^, and 2) Substantial morphological changes that promote the HEs to change shape to make HSPCs^5–8^. During EHT, there is a progressive decrease in the expression of EC markers, such as *kdrl,* and upregulation of hemogenic markers, such as *runx1*^3, 6–8^. EHT-mediated HSPC generation is highly evolutionarily conserved across zebrafish and mammals^9, 10^. In zebrafish, HSPCs form in the ventral wall of the dorsal aorta between 24-48 hours post fertilization (hpf). They then migrate at 2-3 days post fertilization (dpf) to the caudal hematopoietic tissue (CHT) where they undergo expansion, and then differentiate into various blood lineages include thymic T-cells, erythrocytes, macrophages, and neutrophils^3, 11–14^. Transcriptional regulation of this cell transition has been extensively studied identifying RUNX1 and GATA2 as critical transcription factors for EHT^8, 15^. In contrast, the precise RNA processing factors involved in embryonic HSPC formation are less explored.

Splicing of pre-mRNA into mature RNA is a highly orchestrated process that results in a complex and often cell-type selective transcriptome. RNA splicing is categorized by the removal of introns and ligation of exons by the spliceosome^16^ that coordinates both constitutive and alternative splicing (AS), a process that increases proteome diversity by generating a wider array of transcript isoforms^17–20^. Previous studies uncovered a critical function for splicing in HE/HSPC production. Loss of core spliceosomal components, such as Sf3b1 (splicing factor 3b, subunit 1), U2af1, and Sf3a3 negatively impacted embryonic HE and HSPC formation indicating these factors are functionally important in the EHT^21–23^. Moreover, a murine scRNA-seq study revealed that there are dynamic changes to the alternative splicing landscape during EHT that are regulated in part by Serine and arginine rich splicing factor 2 (SRSF2)^24^. Similar to core spliceosomal factors, SRSF2 controls a wide array of splicing decisions, and its loss led to extensive splice isoform changes and disrupted EHT in mice. Coordination by SRSF2 did not explain all EHT alternative splicing decisions identified, suggesting there is much more to learn about its regulation.

While core spliceosomal components are important for global pre-mRNA splicing decisions, cell-type and context-selective alternative splicing is typically modulated by additional factors. For example, *cis*-acting regulatory sequences located within the primary transcript attract *trans-*acting splicing factors such as spliceosome machinery and other sequence-specific RNA binding proteins (RBPs) that modulate splicing decisions^20, 25, 26^. The *cis*-acting regulatory elements can influence whether a transcript is alternatively spliced and the inclusion level in the final transcript^20, 26^. These RBPs can alter splicing by enhancing or repressing inclusion of alternative exons or retained introns ^20, 26, 27^. For example, some factors like SR proteins are splicing enhancers, while others, like HNRNP proteins, repress splicing. There are even factors like polypyrimidine tract binding proteins or STAR proteins, that can either enhance or repress splicing dependent on the location within the transcript where the RBP binds. When binding to Exon/Intron enhancers within the mRNA, these splicing factors often enhance inclusion of the exon by recruiting splicing machinery or blocking recruitment of splicing repressors. Binding of RBPs to exon/intron splicing silencers often represses inclusion of the alternative exon by blocking the recruitment of splicing machinery or blocking recruitment of splicing enhancers^28–30^. For example, Rbfox2 binds to an intronic splicing enhancer downstream of exon 16 of 4.1R encoding EPB41, causing shifts in erythroid differentiation^31, 32^. The identity of the RBPs controlling EHT remain largely unknown.

In this study, using publicly available RNA-seq datasets from EC, HE, and HSPC from different timepoints within the EHT, we elucidated the alternative splicing landscape of the zebrafish EHT and potential *cis*-acting sequence elements enriched in the alternative splice isoforms. We identified several differentially expressed RBPs between ECs and newly forming HE and HSPCs. Through genetic screening of these RBPs, we identified K homology domain containing RNA binding signal transduction associated 1 (Khdrbs1) paralogs as critical regulators involved in the EHT during zebrafish embryogenesis. Khdrbs1, also known as SRC associated in mitosis of 68 kDa (Sam68), is an evolutionarily conserved STAR containing protein which contains one KH domain^33, 34^. Khdrbs1 is ubiquitously expressed in most tissues^35, 36^ and is primarily found in the nucleus^37^. It binds as a homodimer to bipartite RNA sequences as well an interacts with U2AF1 as a stabilizer^38^. Khdrbs1 is known to be involved in a variety of molecular and cellular functions including splicing, alternative polyadenylation, signaling pathways and cell cycle regulation and acts as a poly(A) binding protein^24, 33, 39–42^. In hematopoiesis, KHDRBS1 can regulate *Runx1* polyadenlyation^24^, is phosphorylated in activated T-cells, and regulates T-lymphocytes and T-cell lymphoma in a splicing dependent manner^42–44^. In our study, we uncovered a new function for Khdrbs1 in embryonic HSPC formation. We showed that mutagenesis of both zebrafish KHDRBS1 paralogs, *khdrbs1a* and *khdrbs1b,* leads to decreased HSPC formation, without affecting vascular specification. Downstream of HSPC, loss of *khdrbs1a/b* decreased T-cell and neutrophil numbers while having no effect on macrophages at 6dpf. Analysis of differential splice isoforms between embryonic EC, HE, and HSPC revealed an enrichment for Khdrbs1 binding motifs within all major types of alternative splicing events, implicating it in EHT alternative splicing regulation. In support of Khdrbs1 acting as an EHT splicing modulator, we observed partial restoration of HSPC levels in *khdrbs1a/b* mutagenized embryos upon treatment with the small molecule splicing modulator E7107. Overall, these studies suggest that Khdrbs1 is a crucial RBP for HSPC formation and differentiation in a splicing-mediated mechanism.

## RESULTS

### Defining the alternative splicing landscape of zebrafish EHT

Splicing is an essential process that can modulate cell fate decisions; however little is known which alternative splicing events regulate the EHT. To uncover how splicing might regulate the EHT, we aimed to define the alternative splicing landscape changes occurring during this process in zebrafish embryos. To capture the splicing profile of HE and HSPC, we analyzed published RNA-seq datasets of EC, HE, and HSPC [**Fig 1A**]. The EHT process is a continuum where the transition from EC-HE-HSPC is repeatedly occurring during a given time window of development. In zebrafish, this time is predominantly 24-48 hpf with higher number of HE early and a higher number of HSPCs later. Expression of EC, HE and HSPC marker genes evolve with time with *runx1* primarily marking HE early and a mix of HE/HSPC later^5, 8, 15^. In our analyses, we used 26 hpf HE (*runx1+;kdrl+*) and EC (*kdrl+*) data from Lefkopoulous *et al.*^45^, 36 hpf HE (*runx1+*) and EC (*kdrl+*) data from Li *et al.*^46^, and HSPC (*cd41/itga2b+*) and EC (*kdrl+*) data from Xue *et al.* ^47^. We performed pairwise splicing analysis for each dataset using rMATS to define significantly different splicing events^48^. There are numerous types of alternative splicing events that make up the alternative splicing landscapes [**Fig 1B**]. The most common type is cassette exon splicing or exon skipping in which one alternative exon is flanked by two constitutive exons. Mutually exclusive splicing occurs when there are two alternative exons with only one or the other included in the final transcript ^17–20^. Alternative 5’ splice site (A5SS) and alternative 3’ splice site (A3SS) usage occurs when nearby 5’ or 3’ splice sites at the beginning or end of an intron, respectively, are employed rather than the canonical ones. Retained introns are when an intron is retained in the final transcript.

**Figure 1:**
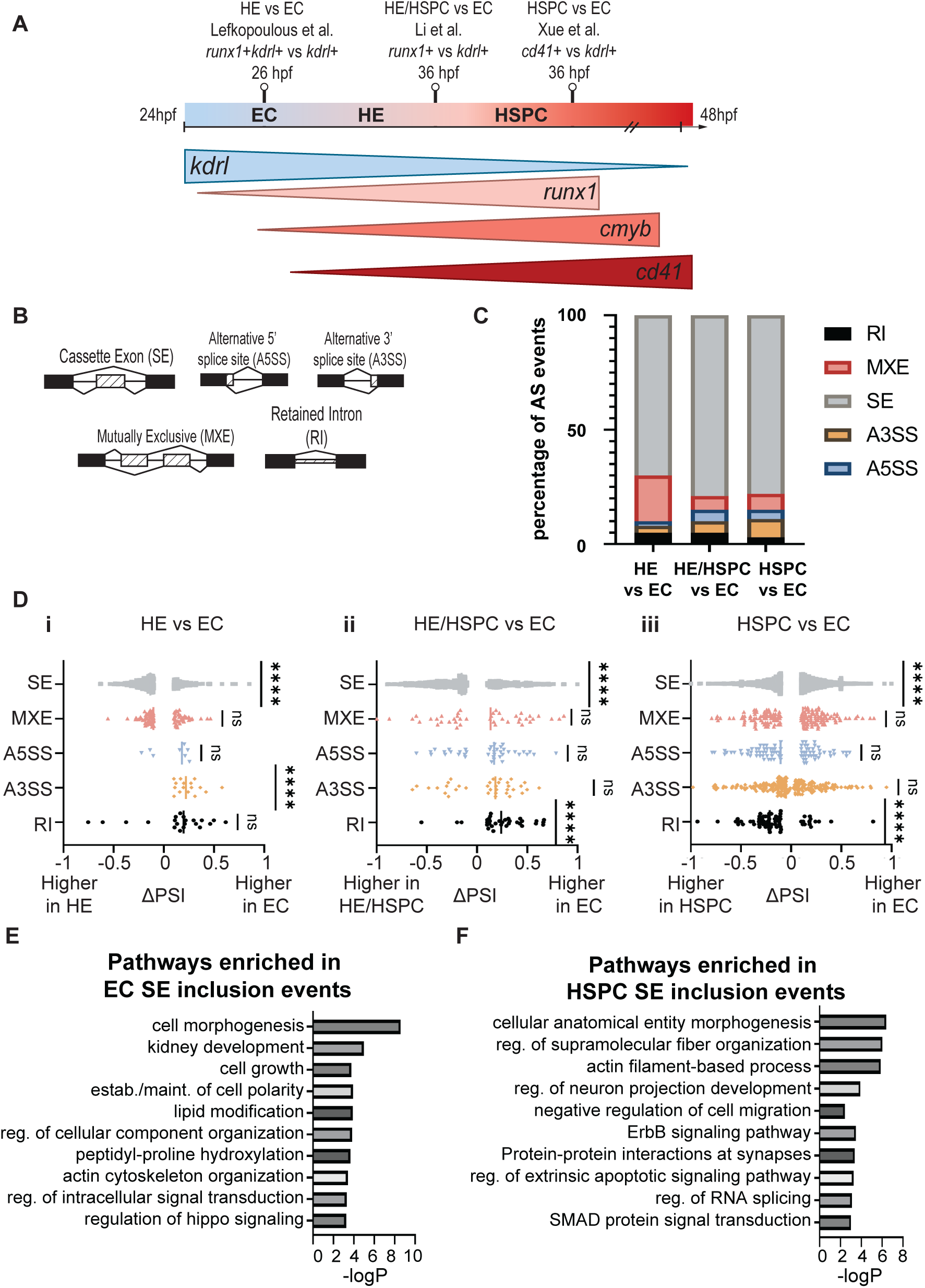
Extensive alternative splicing changes in critical regulators occur during the zebrafish EHT. [**A**] Schematic representing the time points and cell types analyzed for differential splicing analysis on published RNA-seq datasets. The datasets used include cells isolated from zebrafish using *kdrl* to mark ECs. The Lefkopoulos et al^45^ dataset contains ECs and HEs isolated from 26hpf embryos with HEs defined as double positive cells for *runx1* and *kdrl.* The Li et al.^46^ dataset contains ECs and HEs/HSPCs isolated from 36hpf embryos with HE/HSPC defined as positive for *runx1*. The Xue et al^47^ dataset contains ECs and HSPCs isolated from 36hpf embryos with HSPCs defined as positive for *cd41*. [**B**] Schematic representing the types of alternative splicing events analyzed. [**C**] Percentage of each type of alternative splicing event out of the total number of significantly differentially spliced events. Significant values were defined as those with an FDR<0.05 and ΔPSI>10%. [**D**] Global comparisons of significantly differentially spliced events between EC defined by *kdrl* expression to HE defined by *runx1 and kdrl* expression at 26 hpf (i), HE/HSPC defined by *runx1* and *kdrl* expression at 36 hpf (ii), and HSPC defined by *cd41* expression at 36 hpf (iii). Significant events were defined as those with an FDR<0.05 and ΔPSI>10%. Statistical test used was a one-sided t-test; **** represents p-value<0.0001; n.s. not significant. [**E-F**] Graphs denoting the top 10 significantly enrichment pathways in cassette exon events with higher inclusion in ECs [**E**] and HSPCs [**F**] based on analysis of RNA-seq data from Xue et al., 2019^47^.

Across all three datasets, we identified substantial alternative splicing [**Fig 1C-D and Table S1**]. In general, across all three comparisons (HEvsEC, HE/HSPCvsEC, HSPCvsEC) the most abundant alternative splicing event was cassette exon skipping. In the HEvsEC comparison, MXE events were also abundant, while in HE/HSPCvsEC and HSPCvsEC A3SS and A5SS were well represented. Inspection of the magnitude and directionality of the splicing changes further delineated how the splicing landscape evolves through the EHT. For example, there is higher cassette exon inclusion in HE or HE/HSPC compared to EC [**Fig 1D**]. The trend is reversed in the HSPCvsEC comparison with lower cassette exon inclusion in HSPC compared to EC. Similarly, global RI trends show trending towards lower retention in HE, significantly lower in HE/HSPC, but higher retention in HSPC **[Fig 1D]**. The significant shifts in global SE and RI at the transition from HE to HSPC suggests an orchestrated shift in the splicing landscape might be specifically critical for this cellular change. Inline with an important function of these alternative transcripts in the EHT, factors containing cassette exon events with higher inclusion in the ECs compared to HSPCs are enriched in pathways such as “cell growth” and “establishment/maintenance of cell polarity” and those with higher inclusion in HSPCs are enriched in pathways corresponding to morphological changes such as “cellular anatomical entity morphogenesis” and “negative regulation of cellular migration” consistent with the physical changes occurring as HSPC change their shape, bud out of the aorta and enter circulation at the end of the EHT **[Fig 1E-F, Table S2]**. What is not clear is what is regulating these global splicing changes.

### Numerous splicing factors are differentially expressed during zebrafish EHT

RNA binding proteins are proteins that can regulate a variety of molecular processes by binding to different RNA transcripts. One such role is regulation of alternative splicing via interactions with unspliced pre-mRNA transcripts^20, 25, 26^. To identify RBPs that might regulate alternative splicing in the EHT, we examined the expression of RBPs in the Xue *et al.* HSPCvsEC RNA-seq dataset as it is the dataset with the most differential splicing events **[Fig 1C].** We performed differential expression analysis with DESeq2 and identified 3,353 differentially expressed genes, 67 of which are known RBPs recorded in human RBP databases^49, 50^, and 13 of those are known splicing factors **[Fig 2A and Table S3]**. These differentially expressed splicing factors showed relatively consistent expression between replicates and have significant adjusted p-values (< 0.05) and log2fold changes (≥0.32) between EC and HSPC **[Fig 2B-C]**.

**Figure 2:**
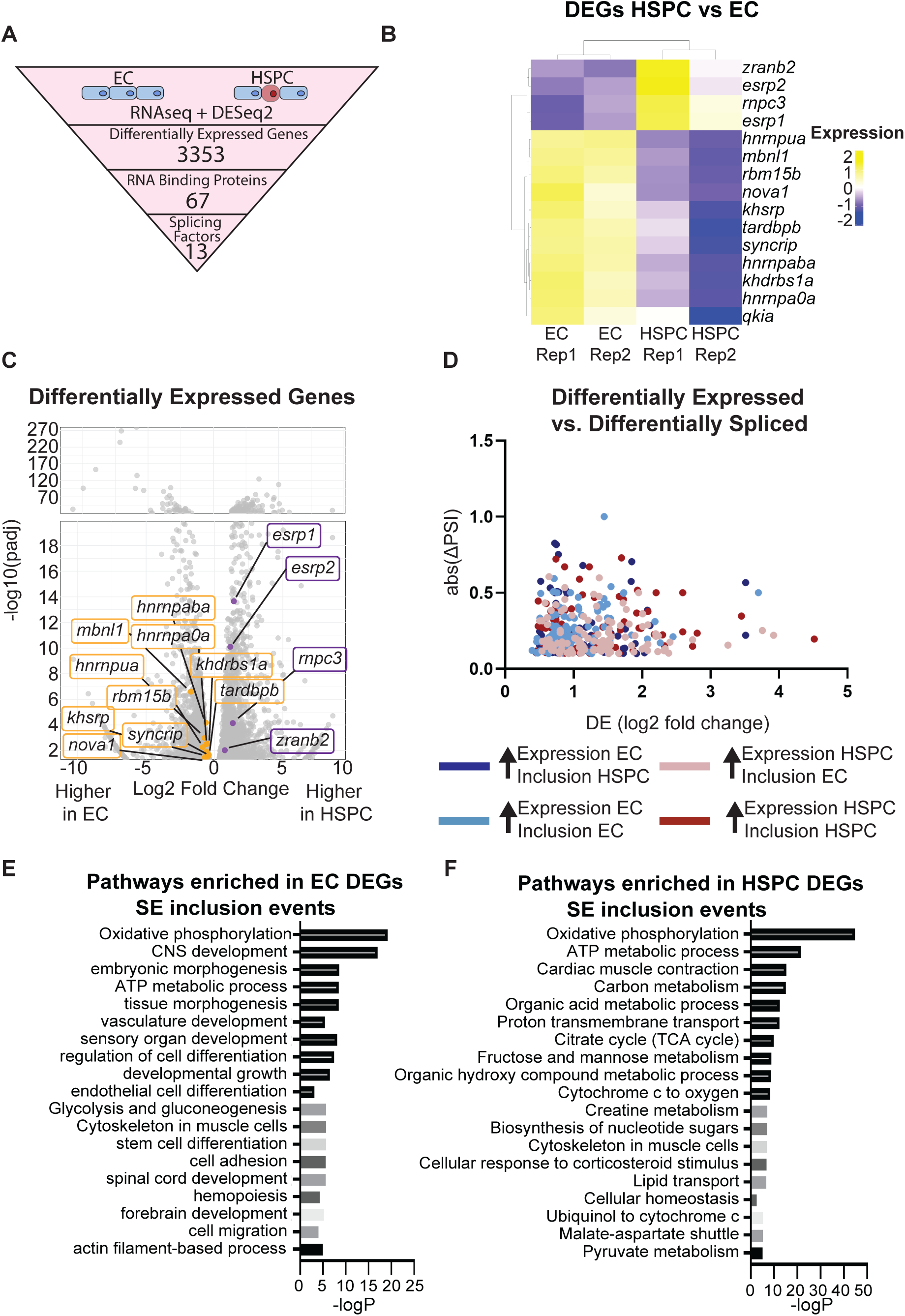
RBP genes are differentially expressed during the zebrafish EHT. [**A**] Schematic representing differential gene expression analysis of RNA-seq data of EC (*kdrl*) and HSPC (*cd41*) from Xue et al., 2019^47^ with a focus on RNA binding protein (RBP) expression. Significant DEG were defined as Log2FoldChange≥0.32 and an adjusted p-value of <0.05. RBPs in the dataset were defined based on RBP2GO^49^ and RBPDB^74^. RBPs with splicing factor functions were defined based on a literature search of all 67 differentially expressed RBPs. [**B**] Heatmap showing normalized differential expression of the 13 differentially expressed splicing factors between EC and HSPC. [**C**] Volcano plot representing log2 fold change vs adjusted p-value of the 3,360 significantly differentially expressed genes. Splicing factors are represented by purple dots while other genes are represented by grey dots. [**D**] Plot representing genes that were both differentially expressed and differentially spliced. Log2 fold change of differential expression graphed against absolute value of ΔPSI. [**E-F**] Graphs denoting pathways enriched in DEGs with higher expression in ECs [**E**] and HSPCs [**F**] using Metascape. One pathway was selected for each summary pathway and graphed by −logP.

Differential splicing can alter the final protein product or result in the introduction of a premature stop codon changing the alternative transcript into a substrate for nonsense-mediated decay, which lowers transcript abundance^51^. Additionally, transcripts expressed significantly low in one cell type compared to the other can appear as false positive alternative splicing^52^. For example, high expression in one cell type might result in more reads covering all possible splice junctions, potentially making it seem like there is differential splicing when the effect is caused by a difference in the number of reads due to higher expression. To determine the impact of differential expression on differential splicing, we compared these two parameters finding little concordance between differential inclusion level and expression differences **[Fig 2D]**. This finding illustrates that the majority of events are likely *bona fide* splicing alterations.

At the pathway level, we did observe convergence in the pathways enriched in alternatively spliced and differentially expressed transcripts **[Fig 2E-F, Table S4]**. Pathways enriched in the DEGs are related to cell processes critical for the EHT, such as “embryonic morphogenesis” and “stem cell differentiation”. Combined our findings illuminate the coordinated changes in the alternative splicing and transcriptional landscape.

### Functional screen identified potential splicing factor regulators of zebrafish EHT

We posited that the differentially expressed splicing factors identified in the transcriptional analysis could be functionally important for the EHT. To test this hypothesis, we screened the impact of mutagenizing the differentially expressed splicing factors on HSPC formation **[Fig 3A]**. We used an F0 CRISPR-Cas9 mutagenesis (aka crispant) approach to induce mutagenesis and then performed *cmyb in situ* hybridization at 36 hpf comparing levels in crispant embryos compared to Cas9 only controls. Those with higher expression in HSPCs were screened individually (*rnpc3* and *zranb2*) or in pairs (*esrp1* and *esrp2*) and those with higher expression in ECs were screened in three pooled groups. Groups were determined based on structural similarities (group #1 all have KH domains), functional similarities (group #3 are heterogeneous nuclear ribonucleoprotein (hnRNP)) proteins, or based on hierarchical clustering (group #2). While *khdrbs1b* was not significantly differentially expressed, we tested it along with its close paralog *khdrbs1a* to account for potential compensation **[Fig 3B]**.

**Figure 3:**
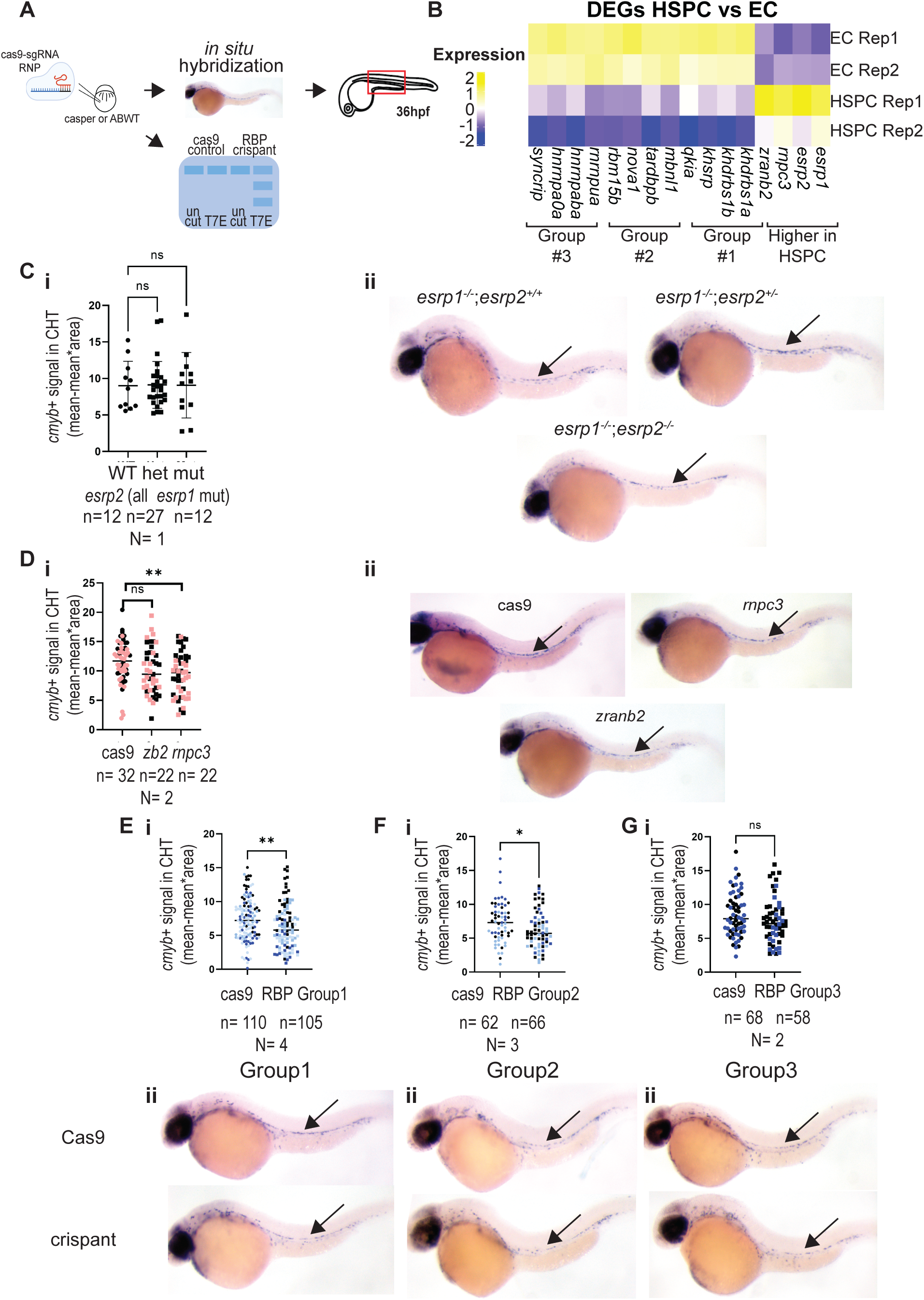
Functional screen of RBPs identified potential regulators of HSPC formation. [**A**] Schema of RBP screening method: Two gRNAs were tested for each gene. The most efficient gRNA for each gene were selected based on a T7 endonuclease mutagenesis assay. For HSPC screening, embryos were injected at 1-4 cell stage with one gRNAs per gene with 3-4 genes tested per pool. HSPC assessment was done using *in situ* hybridization for *cmyb* expression at 36hpf. Mutants or F0 mutagenized embryos (crispants) for splicing factors with higher expression in HSPCs [**C-D**] and ECs [**E-G**] were assessed. Following *in situ* hybridization, embryos were imaged at 8x magnification and pixel intensity within the dorsal aorta area was quantified using image J after 8 bit conversion. [**B**] Heatmap showing the grouping of RBPs used for a pooled CRISPR screen to identify splicing factors functionally involved in HSPC formation. [**C-D**] *cmyb* expression in *esrp1;esrp2* mutants in an AB background [**C**] and in *zranb2* and *rnpc3* crispants in a *cd41:GFP* transgenic background [**D**]. (i) Quantification of *cmyb* expression within the dorsal aorta (DA). Statistical test used for *esrp1;esrp2* was a student t-test with welch’s correction and for *zranb2* and *rnpc3* was one-way ANOVA. ** = pvalue<0.01. Images were analyzed by image J. Different colors represent independent replicates. (ii) Representative images of *cmyb* expression. [**E-G**] *cmyb* expression in Group 1 (*khdrbs1a, khdrbs1b, khsrp, qkia)* crispants in AB and *casper* backgrounds [**E**], Group 2 (*nova1, mbnl1, tardbp, rbm15b)* crispants in the casper background [**F**], and Group 3 (*hnrnpa0a, hnrnpub, hnrnpua, and syncrip)* crispants in AB and *casper* backgrounds [**G**]. (i) Quantification of *cmyb* expression within the DA. All analysis was performed with student t-test with welch’s correction. *pvalue<0.05 and **pvalue<0.01. Different colors represent independent replicates. (ii) Representative images of *cmyb* expression.

Among those with higher expression in HSPCs, double *esrp1* ^-/-^; *esrp2* ^-/-^ mutants and *zranb2* mutagenized embryos had no change in HSPC levels at 36 hpf **[Fig 3C-D]**. In contrast, embryos mutagenized for *rnpc3* had significantly fewer *cmyb+* HSPCs **[Fig 3D]**. Analysis of crispants of the genes with higher inclusion in ECs revealed mutagenesis of those in groups 1 and 2 both caused a decrease in HSPC formation, while group 3 depletion had no impact **[Fig 3E-G]**. This functional screen revealed many possible new players in HSPC formation.

### RBP Khdrbs1a and Khdrbs1b regulate HSPC formation and differentiation

As mutagenesis of the group 1 pool had the most significant and reproducible decrease in *cmyb+* HSPC levels **[Fig 3E]**, we further dissected the effect of these factors. Although mutagenesis of *rnpc3* also showed a significant decrease in HSPCs, we chose not to further explore this factor as it is part of the U12-dependent splicing system and very few of the differentially spliced genes in the EHT analysis are U12 dependent^53^ [data not shown]. To determine which group 1 RBPs contributed to the effect on HSPCs, we tested smaller pools: *qkia+khsrp* and *khdrbs1a+khdrbs1b (kha+b)*. Combined mutagenesis of *qkia* and *khsrp* had no impact on HSPC levels, but *khdrbs1a+b* mutagenesis significantly decreased *cmyb+* HSPC levels **[Fig 4A]**.

**Figure 4:**
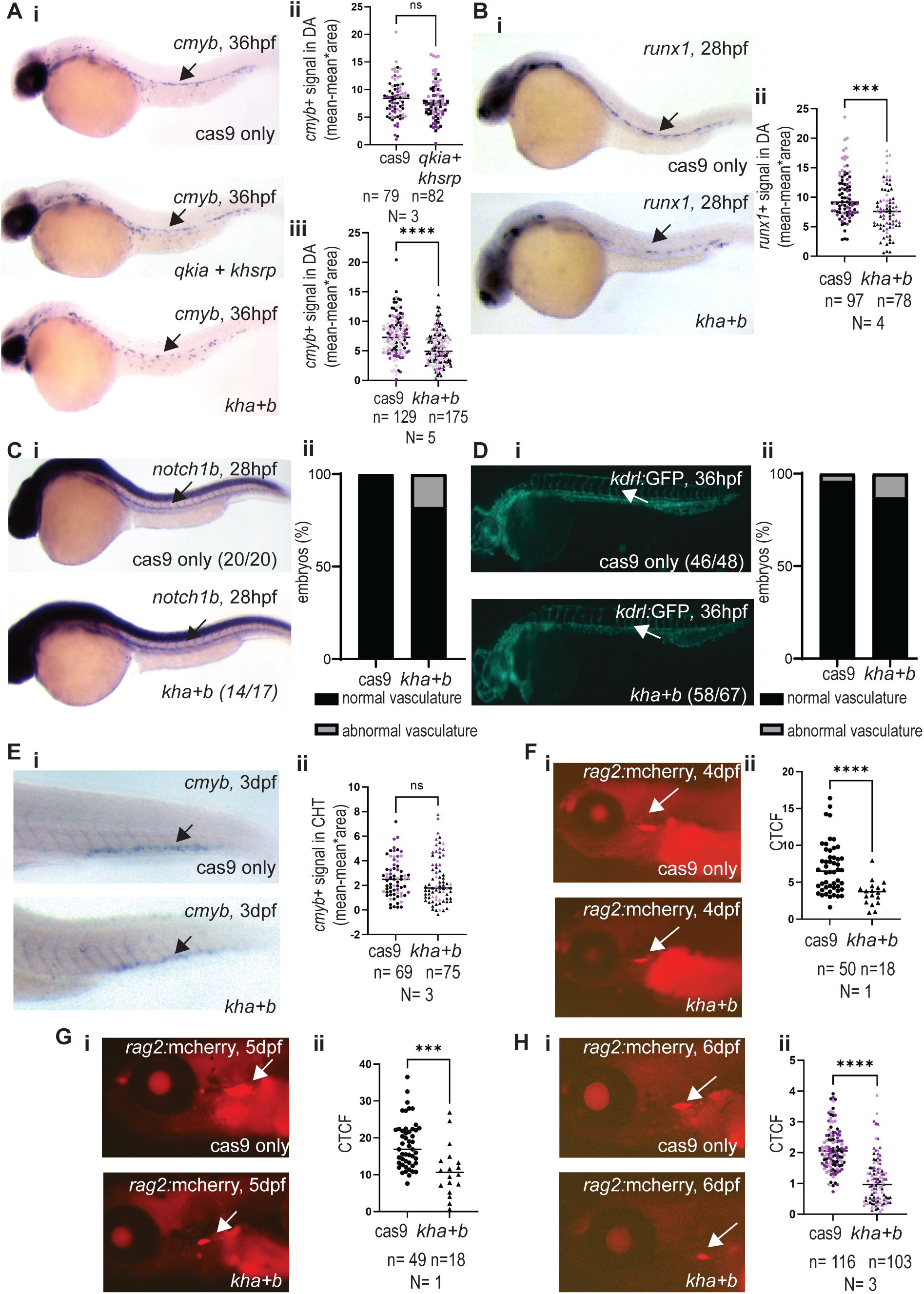
Khdrbs1 regulates HSPC formation and differentiation. [A] HSPC levels in crispants were assessed by *cmyb in situ* hybridization at 36hpf. For each gene tested, one gRNA per RBP was injected for *qkia* and *khsrp* and *khdrbs1a* and *khdrbs1b*. (i) Representative images of *cmyb* expression. (ii-iii) Quantification of *cmyb* expression within the DA in (ii) *qkia+khsrp* and (iii) *khdrbs1a+khdrbs1b (kha+b)* crispants. ****=pvalue<0.0001 calculated by a student t-test with welch’s correction. Images were analyzed by image J. Different colors represent independent replicates. [**B**] HE levels in crispants were assessed by *runx1 in situ* hybridization at 28hpf in *khdrbs1a+b* crispants. (i) Representative images of *runx1* expression. (ii) Quantification of *runx1* expression within the DA. ****=pvalue<0.0001 calculated by student t-test with welch’s correction. Images were analyzed by image J. Different colors represent independent replicates. [**C-D**] EC levels were assessed by *notch1b in situ* hybridization at 28hpf [**C**] and live imaging of *kdrl:*GFP at 36hpf [**D**]. Images were scored by presence and lack of proper vasculature formation. (i) Representative images shown. (ii) Quantification of the percentage of embryos with the designated phenotype. Fisher’s Exact test used. n.s.-not significant. All 28hpf and 36hpf embryos were imaged by dissecting scope at 8x magnification. [**E**] HSPC levels within the caudal hematopoietic tissue (CHT) in crispants were assessed by *cmyb in situ* hybridization at 3dpf. (i) Representative images of *cmyb* expression. (ii) Quantification of *cmyb* expression within the CHT. Images were taken using a dissecting scope at 5x magnification and analyzed by image J. n.s. indicated non-significant. Different colors represent independent replicates. [**F-H**] Live imaging of zebrafish expressing the *rag2*:mcherry transgenes at 4 (**F**), 5 (**G**), and 6dpf (**H**). (i) Representative images of *rag2:mcherry* expression. Larvae were imaged with an inverted scope at 10x magnification [**F-G**] or 5x magnification [**H**]. (ii) Quantification of *rag2:mcherry* expression within the thymus. Intensity of fluorescence in the thymus was quantified in image J. ***pvalue<0.001 and ****pvalue<0.0001 calculated by a student t-test with welch’s correction. Different colors represent independent replicates.

To determine the step in the EHT when Khdrbs1 factors might be acting, we performed analysis of several steps from dorsal aorta formation, HE induction to HSPC formation. Mutagenesis of *khdrbs1a+b* diminished HE levels as evidenced by lower expression of *runx1* within the dorsal aorta and fewer *drl:*mcherry and *kdrl*:GFP double positive cells at 28hpf **[Fig 4B, S1A**. Earlier defects in vascular development can result in catastrophic defects in HE induction and HSPC formation. To determine if the HE and HSPC formation defects in *khdrbs1a+b* crispants were due to vascular defects, we examined *notch1b* expression within the dorsal aorta at 28hpf and *kdrl:GFP* as a marker of all vascular at 36hpf **[Fig 4C-D]**. At both 28hpf and 36hpf, there is no significant defect in the vasculature or dorsal aorta indicating the defect in *khdrbs1a+b* crispant embryos is specifically within the EHT.

After HSPC form, they migrate into the secondary caudal hematopoietic tissue (CHT) where they undergo expansion^3, 54^. To understand the downstream consequence of *khdrbs1a+b* mutagenesis, we examined HSPC levels with the CHT at 3 dpf **[Fig 4E]**. Unlike the decrease of *cmyb+* HSPC within the dorsal aorta at 36 hpf, we see a recovery in the number of HSPCs by 3dpf, indicating that this defect in the EHT does not have long term consequences on HSPC numbers. Next, we wanted to see if there are problems with HSPC quality that impacts downstream differentiation. We examined erythroid, myeloid, and lymphoid blood cells at 6 dpf in *khdrbs1a+b* crispants compared to controls **[Fig 4F-H, S1-2]**. The most striking and reproducible effect of *khdrbs1a+b* mutagenesis was a decrease in *rag1/2+* T-cells at 4dpf, 5dpf, and 6dpf **[Fig 4F-H, S1B-D]**. Mild increases in *gata1:dsRed+* erythroid cells, decreases in *mpx:GFP+* neutrophils, and no change in *mpeg1.1:mCherry+* macrophage numbers were noted also in *khdrbs1a+b* crispants compared to Cas9 only controls **[Fig S2].** Together, these data indicate that Khdrbs1 factors likely affect the differentiation functionality of newly forming HSPCs.

### Khdrbs1 regulates HSPC formation and differentiation via a splicing mechanism

Khdrbs1 has multiple known molecular mechanisms, including impacting alternative polyadenylation, alternative splicing, and acting as a polyA binding protein^24, 33, 40, 42^. To elucidate if the EHT phenotype is splicing mediated, we used an *in silico* approach using RNA MAP Analysis and Protein Server (RMAPS)^55^ to search for any RBP motifs within the differentially spliced events identified between HSPCvsEC. Khdrbs1 was the only differentially expressed splicing factor that showed an enriched binding motif in the sequences of alternative RI and SE splicing events. **[Fig 5A-B, S4]**. Khdrbs1 binding motifs were enriched within the introns upstream and downstream of skipped exons and in the upstream exons of retained introns **[Fig 5A-B]**. This finding corresponds with a significantly decreased splice site strength on the 5’ss of retained introns and 5’ss and 3’ss of skipped exons, consistent^55^ with these being alternative splicing sites **[Fig S4]**.

**Figure 5:**
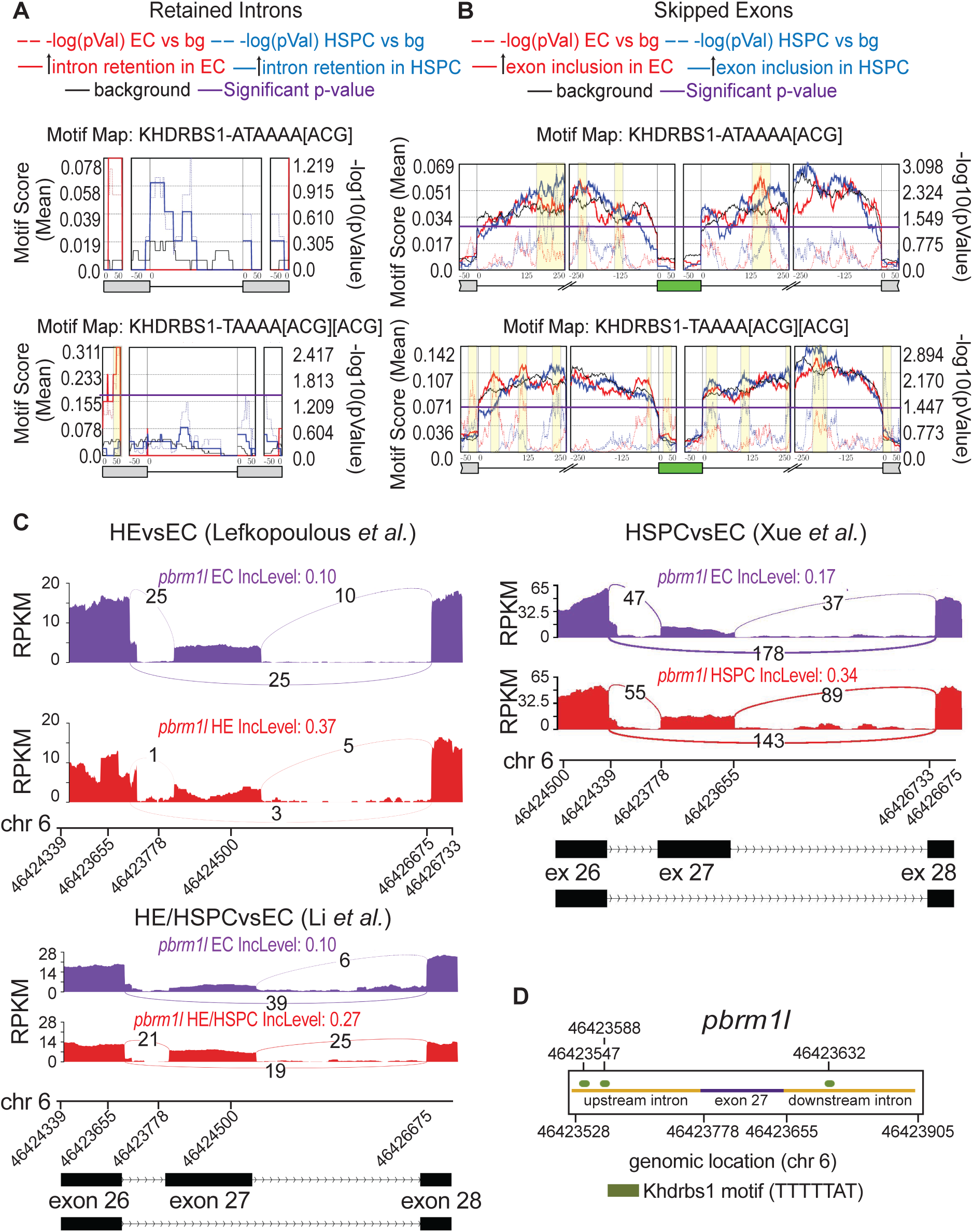
Differential splicing in the zebrafish EHT is potentially regulated by the RBP Khdrbs1. [**A-B**] Metaplot of rMAPS^55^-based Khdrbs1 motif enrichment in significantly differentially spliced RI [**A**] and SE [**B**] events in the HSPCvsEC data from Xue et al., 2019 from Figure 1D (iii). The enrichment of two different Khdrbs1 binding motifs (*ATAAAA[ACG]* and *TAAAA[ACG][ACG]*) were assessed along the entire intervening intron and 50 bp upstream and downstream of the flanking exon junctions for RI events and 50bp +/- exon junctions for SE and adjacent exons plus 250 bp at the start and end of the adjacent introns. Significance cutoff for differentially spliced events is FDR<0.05 and ΔPSI>10%. Solid lines indicate motif score, dotted lines indicated negative log(p-value), and yellow shading indicates significantly enriched peaks. Background events were taken from non-significant values of FDR=1. Significance for the motif enrichment was defined as p-value <0.05 (purple line; −log10(0.05)=1.3). [**C**] Sashimi plot and schematic representation for *pbrm1l* cassette exon 27 and flanking exons 26 and 28 based on rMATS analysis of RNA-seq data Lefkopoulos et al^45^ (HEvsEC), Li et al.^46^ HE/HSPCvsEC, and Xue et al^47^ (HSPCvsEC). [**D**] Schematic representing relative location of Khdrbs1 binding motifs in the flanking introns of the differentially spliced alternative exon 27 in *pbrm1l.* Bases displayed are the exon (purple) and 250 bp upstream and downstream (gold). Location of Khdrbs1 binding sites are represented by dots (green).

We surveyed two KHDRBS1 binding motifs: ATAAAA[ACG] and TAAAA[ACG][ACG]. The bivalent TAAAA[ACG][ACG] motif stronger motif scores and more locations of enrichment along both RI and SE splicing events. An example of a KHDRBS1 containing alternative splicing event is the exon 27 skipped exon in *polybromo1-like* (*pbrm1l*) **[Fig 5C]**. Pbrm1l is the zebrafish ortholog of human PBMR1, a known chromatin remodeler associated with the SWI/SNF complex with established roles in leukemia and HSPC maintenance^56–58^. The alternative exon 27 in *pbrm1l* has higher inclusion in the HE, HE/HSPC, and HSPCs compared to ECs and is one of only two differentially spliced events across the three datasets **[Fig 5C]**. The predicted change in the Pbrm1l proteinwith the alternative exon excluded is a known conserved deletion{Horikawa, 2002 #118} around 55 amino acids near the c-terminus of the protein. This deletion includes a disordered region near the bromodomain and adjacent homology domain 2 that may change flexibility of the protein and impact protein interactions {Morris, 2021 #150}. **[Fig 5D]**.

Based on the higher expression of *khdrbs1a* in ECs compared to HSPCs, the lower inclusion of the SE in ECs, and the enrichment of Khdrbs1 motifs within alternatively spliced transcripts between HE, HSPC, and EC, we predict that Khdrbs1 factors could be acting as splicing repressors within ECs during the EHT. To functionally test this hypothesis, we treated *khdrbs1a+b* crispant embryos with a low dose of the splicing modulator E7107. While this dose of E7107 had no effect on *cmyb+* HSPC levels in the Cas9 only control, it partially restored HSPC to near control levels in *khdrbs1a+b* crispants **[Fig 6]**. This finding was validated in an 8bp-deletion germline mutant for *khdrbs1b* with subsequent *khdrbs1a* mutagenesis **[Fig S5].** Combined the data indicate that Khdrbs1 factors likely regulate HSPC formation through splicing modulation.

**Figure 6:**
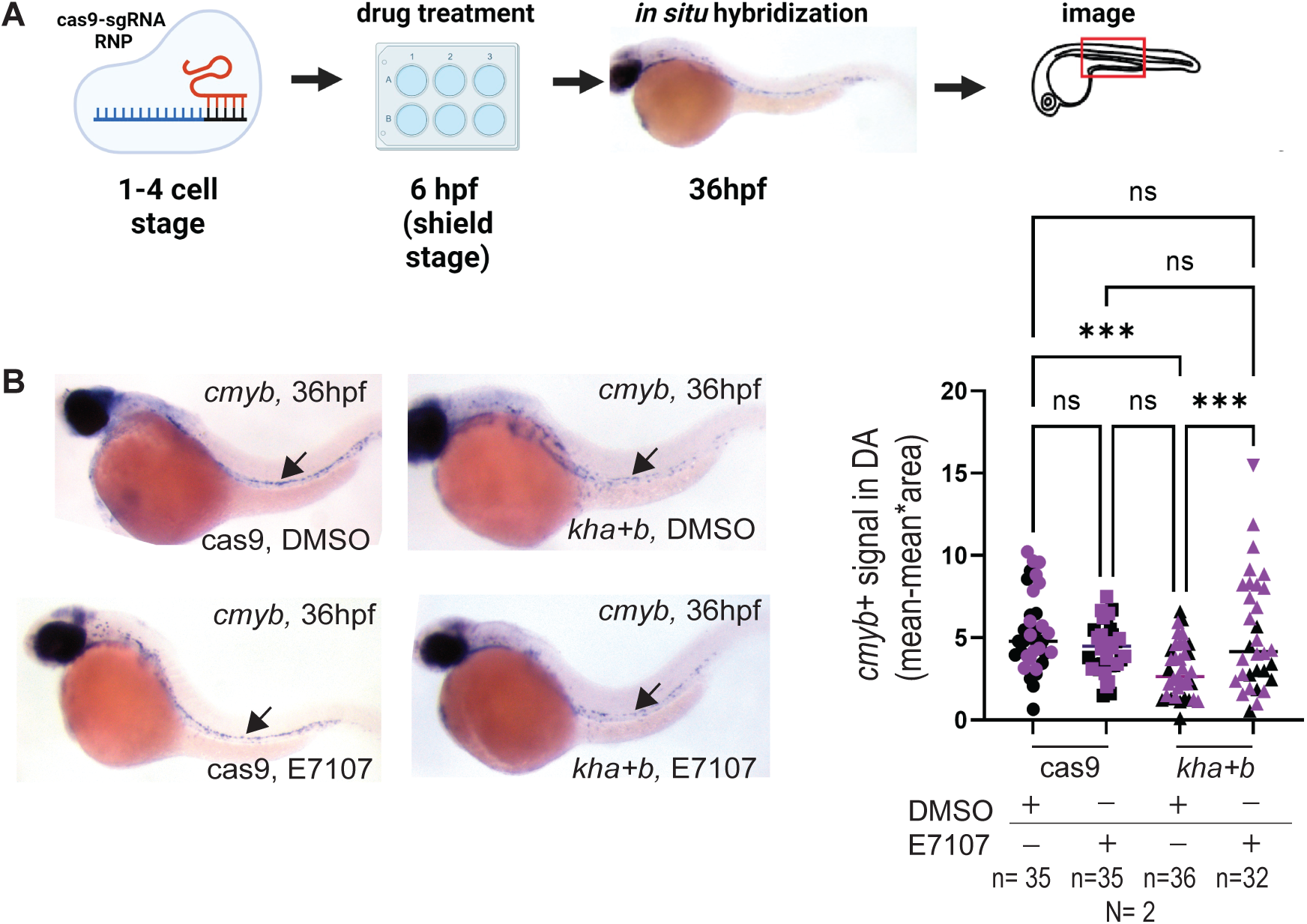
Khdrbs1 regulates HSPC formation via a splicing mechanism. [**A**] Schematic representation of workflow studying the effect of the splicing modulator E7107 on the HSPC formation defects in *khdrbs1a+b* crispants (*kha+b*). Crispants for *khdrbs1a+b* or cas9 only embryos were treated from 6-36hpf with either 30μM E7107 or DMSO. *In situ* hybridization was performed on 36hpf embryos using a *cmyb* probe and then imaged with dissecting scope with 8x magnification. [**B**] Quantification and images of cas9 only and *khdrbs1a+b* crispant embryos at 36hpf with *in situ* hybridization of *cmyb* probe. Images were analyzed by image j after conversion to 8 bit and inversion of colors. Analysis was performed using one-way ANOVA. ** represent p-value <0.001, * represents p-value <0.05 and n.s represents non-significant p-value of >0.05. Different colors represent independent replicates.

## DISCUSSION

In this study, we show that splicing is highly dynamic during the EHT with the differential expression of several RBPs. Through functional assessment, we demonstrate that some of these differentially expressed splicing factors regulate HSPC formation. In particular, we identified Khdrbs1 as a novel regulator of embryonic HSPC formation and differentiation. Its function is selective at the EHT as Khdrbs1 deficiency diminished HE and HSPC formation without impacting vasculature. Although HSPC frequency rebounded after an initial dip in production, Khdrbs1 loss had lasting impacts on HSPC quality with multilineage differentiation defects especially apparent in the lymphoid lineage. These findings identify a novel regulator of HSPC emergence, expanding our current knowledge of molecular drivers of hematopoiesis, and lay the groundwork for potential unexplored interactors and pathways that may regulate embryonic HSPC formation and differentiation.

RBPs that regulate splicing play a crucial role in HSPC emergence. Splicing factors that regulate 3’ss recognition, such as Sf3b1 and U2af1, and their interacting factors, such as Sf3a3, are universally essential for proper splicing. These core splicing factors are known to regulate the EHT, as zebrafish embryos deficient in any one of these factors have a drastic decrease in HSPC formation^21, 22, 59, 60^. As opposed to these core splicing factors like Sf3b1 and U2af1 that are required for recognition of the branch point and polypyrimidine tract^61^, Khdrbs1 is a more selective splicing factor as it regulates specific exons and genes^42^. Our findings uncovered that similar to these core splicing factors, Khdrbs1 modulates embryonic HSPC formation via splicing modulation. The effects downstream of Khdrbs1-regulated splicing important for EHT could be one of a variety of pathways including cell cycle regulation^62, 63^ and signal transduction^44, 64^, such as NFKB^65^ and JAK/STAT3^66^, which are critical for HSPC formation ^59, 67^. The data suggest that Khdrbs1 adds a layer of specificity to the broader RNA regulatory network governing HSPC emergence.

A previous nanopore scRNA-seq study in mice demonstrated substantial differential splicing between aortic endothelial cells, hemogenic endothelial cells and pre-HSCs; however, these data did not differentiate between the different types of alternative splicing events^24^. One limitation of this study is that this long read sequencing does not provide the high depth sequencing coverage required to detect small changes in alternative splicing events that bulk-RNA seq can^68^. Using several bulk RNA-seq datasets from isolated zebrafish EHT cells, we uncovered substantial differential splicing as cells transit from ECs to HSPCs **[Fig 1]**. For example, there is higher cassette exon inclusion in HE or HE/HSPC but lower in HSPC relative to EC. Similarly, global RI changes trend towards lower retention in HE, significantly lower in HE/HSPC, but higher retention in HSPC **[Fig 1D]**. These shifts in global alternative splicing at the transition from HE to HSPC suggests that these large changes to the splicing landscape may be critical for this process. Inline with an important function of these alternative transcripts, we noted enrichment of morphogenesis and migration pathways in the factors containing highly included cassette exons within HSPC, consistent with the utilization of these pathways for cellular egress from within the dorsal aorta. These processes concur with the pathways observed in the previous mouse studies^68^.

Khdrbs1 is known to regulate hematopoietic cells. For example, it is phosphorylated in activated T-cells, regulates alternative splicing in T-lymphocytes and T cell lymphoma, and acts as a regulator of *Runx1* isoform expression in adult HSCs^24, 42–44, 64^. We delineated a new function for Khdrbs1 in embryonic HSPC formation with the underlying molecular mechanism to modulate the EHT occurring in a splicing-dependent manner **[Fig 6]**. In addition to splicing, Khdrbs1 regulates a variety of gene expression steps, such as transcription^69^, alternative polyadenylation^24^ and mRNA stability and export^70^. Although we did not explore these additional functions of Khdrbs1, our data support splicing as at least one of the key mechanism through which Khdrbs1 modulates the EHT.

Khdrbs1 binding motifs were enriched in all major types of alternative splicing events detected between EC and HSPC **[Fig 5 and S3]**. When binding to introns, Khdrbs1 can functions as either an enhancer or a repressor of splicing depending on where it binds within the transcript. When binding to an intronic splicing silencer, Khdrbs1 represses inclusion^71^. It can function as an enhancer when binding to the 3’ss and recruiting U2AF1 for 3’ss recognition, by interacting with HNRNP proteins, or binding to intronic splicing silencers^72, 73^. In alternative splicing events occurring during the EHT transition, we identified an enrichment in Khdrbs1 binding sites on the introns upstream and downstream of SE events and on the upstream exon of RI events, correlating with the observed weaker 3’ss and 5’ss in SE events and only weaker 5’ss in RI events **[Fig S4]**. The data imply that during EHT Khdrbs1 might act as an enhancer to compensate for the weaker splice sites. This seems to be observed in the SE events. In the EC inclusion events, Khdrbs1 binding motifs are enriched in introns upstream of EC-included cassette exons, whereas they are enriched in the downstream introns of SE events included more in HSPCs **[Fig 5]**. Khdrbs1 expression is higher in ECs **[Fig 2]** and could serve to promote exon inclusion of cassette exons. This model is supported by the enrichment of SE included events in ECs compared to HSPCs **[Fig 1]**. In HSPCs, Khdrbs1 motifs are more enriched within introns downstream of cassette exons. This binding position might shift exon inclusion in cells with lower Khdrbs1 levels as HSPCs show more exon exclusion compared to EC. In RI events, our data also support a role for Khdrbs1 act as a splicing enhancer as intron removal is globally better in ECs compared to HSPCs **[Fig 1]**.

While these phenomena seem to be true on a global scale, since Khdrbs1 can act as an enhancer or repressor, Khdrbs1 may function differently for individual events. For example, the upstream and downstream introns of exon 27 in *pbrm1l* contains Khdrbs1 binding motifs **[Fig 5]**. Here, we posit that the sites flanking exon 27 might have opposing roles on exon inclusion and compete for Khdrbs1 binding. The end result is more exon 27 inclusion in HSPC that express lower levels of Khdrbs1. The data illustrate the complexity of splicing regulation through cellular transitions.

Altogether this study defines the splicing landscape of the zebrafish EHT and identifies Khdrbs1 as a novel regulator of the process. We established the role of Khdrbs1 as a previously unknown critical regulator of HSPC formation and identified splicing modulation as a potential mechanism by which Khdrbs1 acts as a regulator in the context of this transition. Our analysis suggests that in the EHT, Khdrbs1 might largely act as a splicing enhancer with transcript-specific functions as a repressor. These studies of EHT-specific splicing regulation in a complex, multicellular context will provide a deeper understanding of how splicing fine-tunes cell fate decisions.

## Supporting information

Karp_SuppFig_Leg

KeyResources

## RESOURCE AVAILABILITY

All resources will be made available by contacting the lead PI.

## ACKNOWLEDGEMENTS

We thank Julie Secombe, Kira Gritsman, Charles Query, Matthew Gamble, and Kristy Stengel for the helpful discussions and support. We thank Varun Gupta for guidance on the computational analysis. We also thank the core facilities at Albert Einstein College of Medicine including the Zebrafish core facility and Flow cytometry core facility (supported by Cancer Center grant [P30CA013330]).

T.V.B. was supported by grants from the National Institutes of Health (NIH) R01DK121738, R01DK131445, and the Edward P. Evans Foundation; I.K. was supported by NIH T32GM007491, NIH F31HL168882, and a Liang Zhu Memorial Fellowship.

## AUTHOR CONTRIBUTIONS

Conceptualization: IK, TVB; Methodology, IK, TVB; Investigation: IK, TVB; Writing—Original Draft: IK; Writing—Review & Editing: TVB; Funding Acquisition: IK, TVB; Supervision: TVB.

## DECLARATION OF GENERATIVE AI AND AI-ASSISTED TECHNOLOGIES

AI was used for assistance with coding. AI was not used for writing of the manuscript.

## SUPPLEMENTAL INFORMATION

Document S1. Figures S1-S4

Table S1: Alternative splicing events defined among HE, HSPC, and EC

Table S2: Pathways enriched in the alternatively spliced events between HE, HSPC and EC

Table S3: Differentially expressed genes between HE, HSPC and EC

Table S4: Pathways enriched in genes differentially expressed between HE, HSPC and EC

Table S5: Sequences of sgRNA and primers used in this study

## REFERENCES

1. Bahary N, Goishi K, Stuckenholz C, Weber G, Leblanc J, Schafer CA, Berman SS, Klagsbrun M, Zon LI. Duplicate VegfA genes and orthologues of the KDR receptor tyrosine kinase family mediate vascular development in the zebrafish. Blood. 2007;110(10):3627–36. Epub 20070814. doi: 10.1182/blood-2006-04-016378. PubMed PMID: 17698971; PMCID: PMC2077312.

2. Nishikawa SI, Nishikawa S, Kawamoto H, Yoshida H, Kizumoto M, Kataoka H, Katsura Y. In vitro generation of lymphohematopoietic cells from endothelial cells purified from murine embryos. Immunity. 1998;8(6):761–9. doi: 10.1016/s1074-7613(00)80581-6. PubMed PMID: 9655490.

3. Kissa K, Herbomel P. Blood stem cells emerge from aortic endothelium by a novel type of cell transition. Nature. 2010;464(7285):112-5. Epub 20100214. doi: 10.1038/nature08761. PubMed PMID: 20154732.

4. Hirschi KK. Hemogenic endothelium during development and beyond. Blood. 2012;119(21):4823–7. Epub 20120313. doi: 10.1182/blood-2011-12-353466. PubMed PMID: 22415753; PMCID: PMC3367889.

5. Ottersbach K. Endothelial-to-haematopoietic transition: an update on the process of making blood. Biochem Soc Trans. 2019;47(2):591–601. Epub 20190322. doi: 10.1042/BST20180320. PubMed PMID: 30902922; PMCID: PMC6490701.

6. Wang Q, Stacy T, Binder M, Marin-Padilla M, Sharpe AH, Speck NA. Disruption of the Cbfa2 gene causes necrosis and hemorrhaging in the central nervous system and blocks definitive hematopoiesis. Proc Natl Acad Sci U S A. 1996;93(8):3444–9. doi: 10.1073/pnas.93.8.3444. PubMed PMID: 8622955; PMCID: PMC39628.

7. Okuda T, van Deursen J, Hiebert SW, Grosveld G, Downing JR. AML1, the target of multiple chromosomal translocations in human leukemia, is essential for normal fetal liver hematopoiesis. Cell. 1996;84(2):321–30. doi: 10.1016/s0092-8674(00)80986-1. PubMed PMID: 8565077.

8. Chen MJ, Yokomizo T, Zeigler BM, Dzierzak E, Speck NA. Runx1 is required for the endothelial to haematopoietic cell transition but not thereafter. Nature. 2009;457(7231):887–91. Epub 20090107. doi: 10.1038/nature07619. PubMed PMID: 19129762; PMCID: PMC2744041.

9. Carroll KJ, North TE. Oceans of opportunity: exploring vertebrate hematopoiesis in zebrafish. Exp Hematol. 2014;42(8):684–96. Epub 20140509. doi: 10.1016/j.exphem.2014.05.002. PubMed PMID: 24816275; PMCID: PMC4461861.

10. Gore AV, Pillay LM, Venero Galanternik M, Weinstein BM. The zebrafish: A fintastic model for hematopoietic development and disease. Wiley Interdiscip Rev Dev Biol. 2018;7(3):e312. Epub 20180213. doi: 10.1002/wdev.312. PubMed PMID: 29436122; PMCID: PMC6785202.

11. Murayama E, Kissa K, Zapata A, Mordelet E, Briolat V, Lin HF, Handin RI, Herbomel P. Tracing hematopoietic precursor migration to successive hematopoietic organs during zebrafish development. Immunity. 2006;25(6):963–75. Epub 20061207. doi: 10.1016/j.immuni.2006.10.015. PubMed PMID: 17157041.

12. Gore AV, Athans B, Iben JR, Johnson K, Russanova V, Castranova D, Pham VN, Butler MG, Williams-Simons L, Nichols JT, Bresciani E, Feldman B, Kimmel CB, Liu PP, Weinstein BM. Epigenetic regulation of hematopoiesis by DNA methylation. Elife. 2016;5:e11813. Epub 20160127. doi: 10.7554/eLife.11813. PubMed PMID: 26814702; PMCID: PMC4744183.

13. Bertrand JY, Chi NC, Santoso B, Teng S, Stainier DY, Traver D. Haematopoietic stem cells derive directly from aortic endothelium during development. Nature. 2010;464(7285):108-11. Epub 20100214. doi: 10.1038/nature08738. PubMed PMID: 20154733; PMCID: PMC2858358.

14. Sood R, English MA, Belele CL, Jin H, Bishop K, Haskins R, McKinney MC, Chahal J, Weinstein BM, Wen Z, Liu PP. Development of multilineage adult hematopoiesis in the zebrafish with a runx1 truncation mutation. Blood. 2010;115(14):2806–9. Epub 20100212. doi: 10.1182/blood-2009-08-236729. PubMed PMID: 20154212; PMCID: PMC2854427.

15. de Pater E, Kaimakis P, Vink CS, Yokomizo T, Yamada-Inagawa T, van der Linden R, Kartalaei PS, Camper SA, Speck N, Dzierzak E. Gata2 is required for HSC generation and survival. J Exp Med. 2013;210(13):2843–50. Epub 20131202. doi: 10.1084/jem.20130751. PubMed PMID: 24297996; PMCID: PMC3865477.

16. Will CL, Luhrmann R. Spliceosome structure and function. Cold Spring Harb Perspect Biol. 2011;3(7). Epub 20110701. doi: 10.1101/cshperspect.a003707. PubMed PMID: 21441581; PMCID: PMC3119917.

17. Wang Y, Liu J, Huang BO, Xu YM, Li J, Huang LF, Lin J, Zhang J, Min QH, Yang WM, Wang XZ. Mechanism of alternative splicing and its regulation. Biomed Rep. 2015;3(2):152–8. Epub 20141217. doi: 10.3892/br.2014.407. PubMed PMID: 25798239; PMCID: PMC4360811.

18. Pan Q, Shai O, Lee LJ, Frey BJ, Blencowe BJ. Deep surveying of alternative splicing complexity in the human transcriptome by high-throughput sequencing. Nat Genet. 2008;40(12):1413–5. Epub 20081102. doi: 10.1038/ng.259. PubMed PMID: 18978789.

19. Wang Z, Burge CB. Splicing regulation: from a parts list of regulatory elements to an integrated splicing code. RNA. 2008;14(5):802–13. Epub 20080327. doi: 10.1261/rna.876308. PubMed PMID: 18369186; PMCID: PMC2327353.

20. Ule J, Blencowe BJ. Alternative Splicing Regulatory Networks: Functions, Mechanisms, and Evolution. Mol Cell. 2019;76(2):329–45. doi: 10.1016/j.molcel.2019.09.017. PubMed PMID: 31626751.

21. De La Garza A, Cameron RC, Nik S, Payne SG, Bowman TV. Spliceosomal component Sf3b1 is essential for hematopoietic differentiation in zebrafish. Exp Hematol. 2016;44(9):826–37 e4. Epub 20160601. doi: 10.1016/j.exphem.2016.05.012. PubMed PMID: 27260753; PMCID: PMC4992596.

22. Burns CE, Galloway JL, Smith AC, Keefe MD, Cashman TJ, Paik EJ, Mayhall EA, Amsterdam AH, Zon LI. A genetic screen in zebrafish defines a hierarchical network of pathways required for hematopoietic stem cell emergence. Blood. 2009;113(23):5776–82. Epub 20090330. doi: 10.1182/blood-2008-12-193607. PubMed PMID: 19332767; PMCID: PMC2700318.

23. Danilova N, Kumagai A, Lin J. p53 upregulation is a frequent response to deficiency of cell-essential genes. PLoS One. 2010;5(12):e15938. Epub 20101231. doi: 10.1371/journal.pone.0015938. PubMed PMID: 21209837; PMCID: PMC3013139.

24. Davis AG, Einstein JM, Zheng D, Jayne ND, Fu XD, Tian B, Yeo GW, Zhang DE. A CRISPR RNA-binding protein screen reveals regulators of RUNX1 isoform generation. Blood Adv. 2021;5(5):1310–23. doi: 10.1182/bloodadvances.2020002090. PubMed PMID: 33656539; PMCID: PMC7948294.

25. De Conti L, Baralle M, Buratti E. Exon and intron definition in pre-mRNA splicing. Wiley Interdiscip Rev RNA. 2013;4(1):49–60. Epub 20121008. doi: 10.1002/wrna.1140. PubMed PMID: 23044818.

26. Fu XD, Ares M, Jr. Context-dependent control of alternative splicing by RNA-binding proteins. Nat Rev Genet. 2014;15(10):689–701. Epub 20140812. doi: 10.1038/nrg3778. PubMed PMID: 25112293; PMCID: PMC4440546.

27. Lunde BM, Moore C, Varani G. RNA-binding proteins: modular design for efficient function. Nat Rev Mol Cell Biol. 2007;8(6):479–90. doi: 10.1038/nrm2178. PubMed PMID: 17473849; PMCID: PMC5507177.

28. Paronetto MP, Achsel T, Massiello A, Chalfant CE, Sette C. The RNA-binding protein Sam68 modulates the alternative splicing of Bcl-x. J Cell Biol. 2007;176(7):929–39. Epub 20070319. doi: 10.1083/jcb.200701005. PubMed PMID: 17371836; PMCID: PMC2064079.

29. Horn T, Gosliga A, Li C, Enculescu M, Legewie S. Position-dependent effects of RNA-binding proteins in the context of co-transcriptional splicing. NPJ Syst Biol Appl. 2023;9(1):1. Epub 20230118. doi: 10.1038/s41540-022-00264-3. PubMed PMID: 36653378; PMCID: PMC9849329.

30. Li D, Yu W, Lai M. Targeting serine- and arginine-rich splicing factors to rectify aberrant alternative splicing. Drug Discov Today. 2023;28(9):103691. Epub 20230627. doi: 10.1016/j.drudis.2023.103691. PubMed PMID: 37385370.

31. Ponthier JL, Schluepen C, Chen W, Lersch RA, Gee SL, Hou VC, Lo AJ, Short SA, Chasis JA, Winkelmann JC, Conboy JG. Fox-2 splicing factor binds to a conserved intron motif to promote inclusion of protein 4.1R alternative exon 16. J Biol Chem. 2006;281(18):12468–74. Epub 20060314. doi: 10.1074/jbc.M511556200. PubMed PMID: 16537540.

32. Quentmeier H, Pommerenke C, Bernhart SH, Dirks WG, Hauer V, Hoffmann S, Nagel S, Siebert R, Uphoff CC, Zaborski M, Drexler HG, Consortium IM-S. RBFOX2 and alternative splicing in B-cell lymphoma. Blood Cancer J. 2018;8(8):77. Epub 20180810. doi: 10.1038/s41408-018-0114-3. PubMed PMID: 30097561; PMCID: PMC6086906.

33. Lukong KE, Richard S. Sam68, the KH domain-containing superSTAR. Biochim Biophys Acta. 2003;1653(2):73–86. doi: 10.1016/j.bbcan.2003.09.001. PubMed PMID: 14643926.

34. Vernet C, Artzt K. STAR, a gene family involved in signal transduction and activation of RNA. Trends Genet. 1997;13(12):479–84. doi: 10.1016/s0168-9525(97)01269-9. PubMed PMID: 9433137.

35. Wang S, Yang Q, Wang Z, Feng S, Li H, Ji D, Zhang S. Evolutionary and Expression Analyses Show Co-option of khdrbs Genes for Origin of Vertebrate Brain. Front Genet. 2017;8:225. Epub 20180104. doi: 10.3389/fgene.2017.00225. PubMed PMID: 29354154; PMCID: PMC5758493.

36. Richard S, Torabi N, Franco GV, Tremblay GA, Chen T, Vogel G, Morel M, Cleroux P, Forget-Richard A, Komarova S, Tremblay ML, Li W, Li A, Gao YJ, Henderson JE. Ablation of the Sam68 RNA binding protein protects mice from age-related bone loss. PLoS Genet. 2005;1(6):e74. Epub 20051216. doi: 10.1371/journal.pgen.0010074. PubMed PMID: 16362077; PMCID: PMC1315279.

37. Derry JJ, Richard S, Valderrama Carvajal H, Ye X, Vasioukhin V, Cochrane AW, Chen T, Tyner AL. Sik (BRK) phosphorylates Sam68 in the nucleus and negatively regulates its RNA binding ability. Mol Cell Biol. 2000;20(16):6114–26. doi: 10.1128/MCB.20.16.6114-6126.2000. PubMed PMID: 10913193; PMCID: PMC86087.

38. Meyer NH, Tripsianes K, Vincendeau M, Madl T, Kateb F, Brack-Werner R, Sattler M. Structural basis for homodimerization of the Src-associated during mitosis, 68-kDa protein (Sam68) Qua1 domain. J Biol Chem. 2010;285(37):28893–901. Epub 20100706. doi: 10.1074/jbc.M110.126185. PubMed PMID: 20610388; PMCID: PMC2937916.

39. Pieraccioli M, Caggiano C, Mignini L, Zhong C, Babini G, Lattanzio R, Di Stasi S, Tian B, Sette C, Bielli P. The transcriptional terminator XRN2 and the RNA-binding protein Sam68 link alternative polyadenylation to cell cycle progression in prostate cancer. Nat Struct Mol Biol. 2022;29(11):1101–12. Epub 20221107. doi: 10.1038/s41594-022-00853-0. PubMed PMID: 36344846; PMCID: PMC9872553.

40. Rajan P, Gaughan L, Dalgliesh C, El-Sherif A, Robson CN, Leung HY, Elliott DJ. The RNA-binding and adaptor protein Sam68 modulates signal-dependent splicing and transcriptional activity of the androgen receptor. J Pathol. 2008;215(1):67–77. doi: 10.1002/path.2324. PubMed PMID: 18273831.

41. Frisone P, Pradella D, Di Matteo A, Belloni E, Ghigna C, Paronetto MP. SAM68: Signal Transduction and RNA Metabolism in Human Cancer. Biomed Res Int. 2015;2015:528954. Epub 20150726. doi: 10.1155/2015/528954. PubMed PMID: 26273626; PMCID: PMC4529925.

42. Matter N, Herrlich P, Konig H. Signal-dependent regulation of splicing via phosphorylation of Sam68. Nature. 2002;420(6916):691-5. doi: 10.1038/nature01153. PubMed PMID: 12478298.

43. Debackere K, Marcelis L, Demeyer S, Vanden Bempt M, Mentens N, Gielen O, Jacobs K, Broux M, Verhoef G, Michaux L, Graux C, Wlodarska I, Gaulard P, de Leval L, Tousseyn T, Cools J, Dierickx D. Fusion transcripts FYN-TRAF3IP2 and KHDRBS1-LCK hijack T cell receptor signaling in peripheral T-cell lymphoma, not otherwise specified. Nat Commun. 2021;12(1):3705. Epub 20210617. doi: 10.1038/s41467-021-24037-4. PubMed PMID: 34140493; PMCID: PMC8211700.

44. Fusaki N, Iwamatsu A, Iwashima M, Fujisawa J. Interaction between Sam68 and Src family tyrosine kinases, Fyn and Lck, in T cell receptor signaling. Journal of Biological Chemistry. 1997;272(10):6214–9. doi: DOI 10.1074/jbc.272.10.6214. PubMed PMID: WOS:A1997WM64700020.

45. Lefkopoulos S, Polyzou A, Derecka M, Bergo V, Clapes T, Cauchy P, Jerez-Longres C, Onishi-Seebacher M, Yin N, Martagon-Calderón N-A, Potts KS, Klaeylé L, Liu F, Bowman TV, Jenuwein T, Mione MC, Trompouki E. Repetitive Elements Trigger RIG-I-like Receptor Signaling that Regulates the Emergence of Hematopoietic Stem and Progenitor Cells. Immunity. 2020;53(5):934–51.e9. doi: 10.1016/j.immuni.2020.10.007.

46. Li Y, Tang C, Liu F, Zhu C, Liu F, Zhu P, Wang L. DNA methylation safeguards the generation of hematopoietic stem and progenitor cells by repression of Notch signaling. Development. 2022;149(10). Epub 20220525. doi: 10.1242/dev.200390. PubMed PMID: 35502759; PMCID: PMC9188753.

47. Xue Y, Liu D, Cui G, Ding Y, Ai D, Gao S, Zhang Y, Suo S, Wang X, Lv P, Zhou C, Li Y, Chen X, Peng G, Jing N, Han J-DJ, Liu F. A 3D Atlas of Hematopoietic Stem and Progenitor Cell Expansion by Multi-dimensional RNA-Seq Analysis. Cell Reports. 2019;27(5):1567–78.e5. doi: 10.1016/j.celrep.2019.04.030.

48. Shen S, Park JW, Lu ZX, Lin L, Henry MD, Wu YN, Zhou Q, Xing Y. rMATS: robust and flexible detection of differential alternative splicing from replicate RNA-Seq data. Proc Natl Acad Sci U S A. 2014;111(51):E5593–601. Epub 20141205. doi: 10.1073/pnas.1419161111. PubMed PMID: 25480548; PMCID: PMC4280593.

49. Caudron-Herger M, Jansen RE, Wassmer E, Diederichs S. RBP2GO: a comprehensive pan-species database on RNA-binding proteins, their interactions and functions. Nucleic Acids Research. 2021;49(D1):D425–D36. doi: 10.1093/nar/gkaa1040.

50. Cook KB, Kazan H, Zuberi K, Morris Q, Hughes TR. RBPDB: a database of RNA-binding specificities. Nucleic Acids Res. 2011;39(Database issue):D301–8. Epub 20101029. doi: 10.1093/nar/gkq1069. PubMed PMID: 21036867; PMCID: PMC3013675.

51. Lewis BP, Green RE, Brenner SE. Evidence for the widespread coupling of alternative splicing and nonsense-mediated mRNA decay in humans. Proc Natl Acad Sci U S A. 2003;100(1):189–92. Epub 20021226. doi: 10.1073/pnas.0136770100. PubMed PMID: 12502788; PMCID: PMC140922.

52. Welch JD, Hu Y, Prins JF. Robust detection of alternative splicing in a population of single cells. Nucleic Acids Res. 2016;44(8):e73. Epub 20160105. doi: 10.1093/nar/gkv1525. PubMed PMID: 26740580; PMCID: PMC4856971.

53. Benecke H, Lührmann R, Will CL. The U11/U12 snRNP 65K protein acts as a molecular bridge, binding the U12 snRNA and U11-59K protein. Embo J. 2005;24(17):3057–69. doi: 10.1038/sj.emboj.7600765. PubMed PMID: WOS:000231789700010.

54. Jin SW, Beis D, Mitchell T, Chen JN, Stainier DY. Cellular and molecular analyses of vascular tube and lumen formation in zebrafish. Development. 2005;132(23):5199–209. Epub 20051026. doi: 10.1242/dev.02087. PubMed PMID: 16251212.

55. Park JW, Jung S, Rouchka EC, Tseng YT, Xing Y. rMAPS: RNA map analysis and plotting server for alternative exon regulation. Nucleic Acids Res. 2016;44(W1):W333–8. Epub 20160512. doi: 10.1093/nar/gkw410. PubMed PMID: 27174931; PMCID: PMC4987942.

56. Nargund AM, Pham CG, Dong Y, Wang PI, Osmangeyoglu HU, Xie Y, Aras O, Han S, Oyama T, Takeda S, Ray CE, Dong Z, Berge M, Hakimi AA, Monette S, Lekaye CL, Koutcher JA, Leslie CS, Creighton CJ, Weinhold N, Lee W, Tickoo SK, Wang Z, Cheng EH, Hsieh JJ. The SWI/SNF Protein PBRM1 Restrains VHL-Loss-Driven Clear Cell Renal Cell Carcinoma. Cell Rep. 2017;18(12):2893–906. doi: 10.1016/j.celrep.2017.02.074. PubMed PMID: 28329682; PMCID: PMC5431084.

57. Li BE, Li GY, Cai W, Zhu Q, Seruggia D, Fujiwara Y, Vakoc CR, Orkin SH. In vivo CRISPR/Cas9 screening identifies Pbrm1 as a regulator of myeloid leukemia development in mice. Blood Adv. 2023;7(18):5281–93. doi: 10.1182/bloodadvances.2022009455. PubMed PMID: 37428871; PMCID: PMC10506108.

58. Wanior M, Kramer A, Knapp S, Joerger AC. Exploiting vulnerabilities of SWI/SNF chromatin remodelling complexes for cancer therapy. Oncogene. 2021;40(21):3637–54. Epub 20210503. doi: 10.1038/s41388-021-01781-x. PubMed PMID: 33941852; PMCID: PMC8154588.

59. Potts KS, Cameron RC, Metidji A, Ghazale N, Wallace L, Leal-Cervantes AI, Palumbo R, Barajas JM, Gupta V, Aluri S, Pradhan K, Myers JA, McKinstry M, Bai X, Choudhary GS, Shastri A, Verma A, Obeng EA, Bowman TV. Splicing factor deficits render hematopoietic stem and progenitor cells sensitive to STAT3 inhibition. Cell Rep. 2022;41(11):111825. doi: 10.1016/j.celrep.2022.111825. PubMed PMID: 36516770; PMCID: PMC9994853.

60. Danilova N, Kumagai A, Lin J. p53 Upregulation Is a Frequent Response to Deficiency of Cell-Essential Genes. Plos One. 2010;5(12). doi: ARTN e15938 10.1371/journal.pone.0015938. PubMed PMID: WOS:000285838900051.

61. Gozani O, Potashkin J, Reed R. A potential role for U2AF-SAP 155 interactions in recruiting U2 snRNP to the branch site. Mol Cell Biol. 1998;18(8):4752–60. doi: 10.1128/MCB.18.8.4752. PubMed PMID: 9671485; PMCID: PMC109061.

62. Li QH, Haga I, Shimizu T, Itoh M, Kurosaki T, Fujisawa J. Retardation of the G2-M phase progression on gene disruption of RNA binding protein Sam68 in the DT40 cell line. FEBS Lett. 2002;525(1-3):145–50. doi: 10.1016/s0014-5793(02)03103-4. PubMed PMID: 12163178.

63. Barlat I, Maurier F, Duchesne M, Guitard E, Tocque B, Schweighoffer F. A role for Sam68 in cell cycle progression antagonized by a spliced variant within the KH domain. Journal of Biological Chemistry. 1997;272(6):3129–32. doi: DOI 10.1074/jbc.272.6.3129. PubMed PMID: WOS:A1997WG19200002.

64. Lang V, Mege D, Semichon M, Gary-Gouy H, Bismuth G. A dual participation of ZAP-70 and scr protein tyrosine kinases is required for TCR-induced tyrosine phosphorylation of Sam68 in Jurkat T cells. Eur J Immunol. 1997;27(12):3360–7. doi: 10.1002/eji.1830271235. PubMed PMID: 9464824.

65. Fu K, Sun X, Wier EM, Hodgson A, Liu Y, Sears CL, Wan F. Sam68/KHDRBS1 is critical for colon tumorigenesis by regulating genotoxic stress-induced NF-kappaB activation. Elife. 2016;5. Epub 20160725. doi: 10.7554/eLife.15018. PubMed PMID: 27458801; PMCID: PMC4959885.

66. Bastidas Torres AN, Cats D, Out-Luiting JJ, Fanoni D, Mei H, Venegoni L, Willemze R, Vermeer MH, Berti E, Tensen CP. Deregulation of JAK2 signaling underlies primary cutaneous CD8(+) aggressive epidermotropic cytotoxic T-cell lymphoma. Haematologica. 2022;107(3):702–14. Epub 20220301. doi: 10.3324/haematol.2020.274506. PubMed PMID: 33792220; PMCID: PMC8883537.

67. Espin-Palazon R, Stachura DL, Campbell CA, Garcia-Moreno D, Del Cid N, Kim AD, Candel S, Meseguer J, Mulero V, Traver D. Proinflammatory signaling regulates hematopoietic stem cell emergence. Cell. 2014;159(5):1070–85. Epub 20141106. doi: 10.1016/j.cell.2014.10.031. PubMed PMID: 25416946; PMCID: PMC4243083.

68. Wang F, Tan P, Zhang P, Ren Y, Zhou J, Li Y, Hou S, Li S, Zhang L, Ma Y, Wang C, Tang W, Wang X, Huo Y, Hu Y, Cui T, Niu C, Wang D, Liu B, Lan Y, Yu J. Single-cell architecture and functional requirement of alternative splicing during hematopoietic stem cell formation. Sci Adv. 2022;8(1):eabg5369. Epub 20220107. doi: 10.1126/sciadv.abg5369. PubMed PMID: 34995116; PMCID: PMC8741192.

69. Yang JP, Reddy TR, Truong KT, Suhasini M, Wong-Staal F. Functional interaction of Sam68 and heterogeneous nuclear ribonucleoprotein K. Oncogene. 2002;21(47):7187–94. doi: 10.1038/sj.onc.1205759. PubMed PMID: 12370808.

70. Wong TL, Loh JJ, Lu S, Yan HHN, Siu HC, Xi R, Chan D, Kam MJF, Zhou L, Tong M, Copland JA, Chen L, Yun JP, Leung SY, Ma S. ADAR1-mediated RNA editing of SCD1 drives drug resistance and self-renewal in gastric cancer. Nat Commun. 2023;14(1):2861. Epub 20230519. doi: 10.1038/s41467-023-38581-8. PubMed PMID: 37208334; PMCID: PMC10199093.

71. Pedrotti S, Bielli P, Paronetto MP, Ciccosanti F, Fimia GM, Stamm S, Manley JL, Sette C. The splicing regulator Sam68 binds to a novel exonic splicing silencer and functions in SMN2 alternative splicing in spinal muscular atrophy. Embo J. 2010;29(7):1235–47. Epub 20100225. doi: 10.1038/emboj.2010.19. PubMed PMID: 20186123; PMCID: PMC2857462.

72. Nadal M, Anton R, Dorca-Arevalo J, Estebanez-Perpina E, Tizzano EF, Fuentes-Prior P. Structure and function analysis of Sam68 and hnRNP A1 synergy in the exclusion of exon 7 from SMN2 transcripts. Protein Sci. 2023;32(4):e4553. doi: 10.1002/pro.4553. PubMed PMID: 36560896; PMCID: PMC10031812.

73. Tisserant A, Konig H. Signal-regulated Pre-mRNA occupancy by the general splicing factor U2AF. PLoS One. 2008;3(1):e1418. Epub 20080109. doi: 10.1371/journal.pone.0001418. PubMed PMID: 18183298; PMCID: PMC2169300.

74. Cook KB, Kazan H, Zuberi K, Morris Q, Hughes TR. RBPDB: a database of RNA-binding specificities. Nucleic Acids Research. 2010;39(Database):D301–D8. doi: 10.1093/nar/gkq1069.

75. Lawrence C. Advances in zebrafish husbandry and management. Methods Cell Biol. 2011;104:429–51. doi: 10.1016/B978-0-12-374814-0.00023-9. PubMed PMID: 21924176.

76. Mathias JR, Perrin BJ, Liu TX, Kanki J, Look AT, Huttenlocher A. Resolution of inflammation by retrograde chemotaxis of neutrophils in transgenic zebrafish. J Leukoc Biol. 2006;80(6):1281–8. Epub 20060908. doi: 10.1189/jlb.0506346. PubMed PMID: 16963624.

77. Bojarczuk A, Miller KA, Hotham R, Lewis A, Ogryzko NV, Kamuyango AA, Frost H, Gibson RH, Stillman E, May RC, Renshaw SA, Johnston SA. Cryptococcus neoformans Intracellular Proliferation and Capsule Size Determines Early Macrophage Control of Infection. Sci Rep. 2016;6:21489. Epub 20160218. doi: 10.1038/srep21489. PubMed PMID: 26887656; PMCID: PMC4757829.

78. Seiler C, Gebhart N, Zhang Y, Shinton SA, Li YS, Ross NL, Liu X, Li Q, Bilbee AN, Varshney GK, LaFave MC, Burgess SM, Balciuniene J, Balciunas D, Hardy RR, Kappes DJ, Wiest DL, Rhodes J. Mutagenesis Screen Identifies agtpbp1 and eps15L1 as Essential for T lymphocyte Development in Zebrafish. PLoS One. 2015;10(7):e0131908. Epub 20150710. doi: 10.1371/journal.pone.0131908. PubMed PMID: 26161877; PMCID: PMC4498767.

79. Carroll SH, Macias Trevino C, Li EB, Kawasaki K, Myers N, Hallett SA, Alhazmi N, Cotney J, Carstens RP, Liao EC. An Irf6-Esrp1/2 regulatory axis controls midface morphogenesis in vertebrates. Development. 2020;147(24). Epub 20201223. doi: 10.1242/dev.194498. PubMed PMID: 33234718; PMCID: PMC7774891.

80. Kimmel CB, Ballard WW, Kimmel SR, Ullmann B, Schilling TF. Stages of embryonic development of the zebrafish. Dev Dyn. 1995;203(3):253–310. doi: 10.1002/aja.1002030302. PubMed PMID: 8589427.

81. Lawson ND, Scheer N, Pham VN, Kim CH, Chitnis AB, Campos-Ortega JA, Weinstein BM. Notch signaling is required for arterial-venous differentiation during embryonic vascular development. Development. 2001;128(19):3675–83. PubMed PMID: WOS:000171974300003.

82. Burns CE, DeBlasio T, Zhou Y, Zhang J, Zon L, Nimer SD. Isolation and characterization of runxa and runxb, zebrafish members of the runt family of transcriptional regulators. Exp Hematol. 2002;30(12):1381–9. doi: 10.1016/s0301-472x(02)00955-4. PubMed PMID: 12482499.

83. Liao EC, Paw BH, Oates AC, Pratt SJ, Postlethwait JH, Zon LI. SCL/Tal-1 transcription factor acts downstream of cloche to specify hematopoietic and vascular progenitors in zebrafish. Genes Dev. 1998;12(5):621–6. doi: 10.1101/gad.12.5.621. PubMed PMID: 9499398; PMCID: PMC316577.

84. Jessen JR, Willett CE, Lin S. Artificial chromosome transgenesis reveals long-distance negative regulation of rag1 in zebrafish. Nat Genet. 1999;23(1):15–6. doi: 10.1038/12609. PubMed PMID: 10471489.

85. Thisse C, Thisse B. High-resolution in situ hybridization to whole-mount zebrafish embryos. Nat Protoc. 2008;3(1):59–69. doi: 10.1038/nprot.2007.514. PubMed PMID: 18193022.

86. Schneider CA, Rasband WS, Eliceiri KW. NIH Image to ImageJ: 25 years of image analysis. Nature Methods. 2012;9(7):671–5. doi: 10.1038/nmeth.2089.

87. Dobin A, Davis CA, Schlesinger F, Drenkow J, Zaleski C, Jha S, Batut P, Chaisson M, Gingeras TR. STAR: ultrafast universal RNA-seq aligner. Bioinformatics. 2013;29(1):15–21. Epub 20121025. doi: 10.1093/bioinformatics/bts635. PubMed PMID: 23104886; PMCID: PMC3530905.

88. Liao Y, Smyth GK, Shi W. featureCounts: an efficient general purpose program for assigning sequence reads to genomic features. Bioinformatics. 2014;30(7):923–30. Epub 20131113. doi: 10.1093/bioinformatics/btt656. PubMed PMID: 24227677.

89. Anders S, Huber W. Differential expression analysis for sequence count data. Genome Biol. 2010;11(10):R106. Epub 20101027. doi: 10.1186/gb-2010-11-10-r106. PubMed PMID: 20979621; PMCID: PMC3218662.

90. BioMart – biological queries made easy.

91. Smedley D, Haider S, Ballester B, Holland R, London D, Thorisson G, Kasprzyk A. BioMart--biological queries made easy. BMC Genomics. 2009;10:22. Epub 20090114. doi: 10.1186/1471-2164-10-22. PubMed PMID: 19144180; PMCID: PMC2649164.

92. Zhou Y, Zhou B, Pache L, Chang M, Khodabakhshi AH, Tanaseichuk O, Benner C, Chanda SK. Metascape provides a biologist-oriented resource for the analysis of systems-level datasets. Nat Commun. 2019;10(1):1523. Epub 20190403. doi: 10.1038/s41467-019-09234-6. PubMed PMID: 30944313; PMCID: PMC6447622.

93. Hwang WY, Fu Y, Reyon D, Maeder ML, Tsai SQ, Sander JD, Peterson RT, Yeh JR, Joung JK. Efficient genome editing in zebrafish using a CRISPR-Cas system. Nat Biotechnol. 2013;31(3):227–9. Epub 20130129. doi: 10.1038/nbt.2501. PubMed PMID: 23360964; PMCID: PMC3686313.

94. Sentmanat MF, Peters ST, Florian CP, Connelly JP, Pruett-Miller SM. A Survey of Validation Strategies for CRISPR-Cas9 Editing. Sci Rep. 2018;8(1):888. Epub 20180117. doi: 10.1038/s41598-018-19441-8. PubMed PMID: 29343825; PMCID: PMC5772360.

95. Labun K, Montague TG, Gagnon JA, Thyme SB, Valen E. CHOPCHOP v2: a web tool for the next generation of CRISPR genome engineering. Nucleic Acids Res. 2016;44(W1):W272–6. Epub 20160516. doi: 10.1093/nar/gkw398. PubMed PMID: 27185894; PMCID: PMC4987937.

96. Labun K, Montague TG, Krause M, Torres Cleuren YN, Tjeldnes H, Valen E. CHOPCHOP v3: expanding the CRISPR web toolbox beyond genome editing. Nucleic Acids Res. 2019;47(W1):W171–W4. doi: 10.1093/nar/gkz365. PubMed PMID: 31106371; PMCID: PMC6602426.

97. Montague TG, Cruz JM, Gagnon JA, Church GM, Valen E. CHOPCHOP: a CRISPR/Cas9 and TALEN web tool for genome editing. Nucleic Acids Res. 2014;42(Web Server issue):W401–7. Epub 20140526. doi: 10.1093/nar/gku410. PubMed PMID: 24861617; PMCID: PMC4086086.

98. Grant CE, Bailey TL, Noble WS. FIMO: scanning for occurrences of a given motif. Bioinformatics. 2011;27(7):1017–8. Epub 20110216. doi: 10.1093/bioinformatics/btr064. PubMed PMID: 21330290; PMCID: PMC3065696.

99. Yeo G, Burge CB. Maximum entropy modeling of short sequence motifs with applications to RNA splicing signals. J Comput Biol. 2004;11(2-3):377–94. doi: 10.1089/1066527041410418. PubMed PMID: 15285897.

100. Xue Y, Liu D, Cui G, Ding Y, Ai D, Gao S, Zhang Y, Suo S, Wang X, Lv P, Zhou C, Li Y, Chen X, Peng G, Jing N, Han JJ, Liu F. A 3D Atlas of Hematopoietic Stem and Progenitor Cell Expansion by Multi-dimensional RNA-Seq Analysis. Cell Rep. 2019;27(5):1567–78 e5. doi: 10.1016/j.celrep.2019.04.030. PubMed PMID: 31042481.

101. Renshaw SA, Loynes CA, Trushell DM, Elworthy S, Ingham PW, Whyte MK. A transgenic zebrafish model of neutrophilic inflammation. Blood. 2006;108(13):3976–8. Epub 20060822. doi: 10.1182/blood-2006-05-024075. PubMed PMID: 16926288.

102. Ellett F, Pase L, Hayman JW, Andrianopoulos A, Lieschke GJ. mpeg1 promoter transgenes direct macrophage-lineage expression in zebrafish. Blood. 2011;117(4):e49–56. Epub 20101117. doi: 10.1182/blood-2010-10-314120. PubMed PMID: 21084707; PMCID: PMC3056479.

103. Traver D, Paw BH, Poss KD, Penberthy WT, Lin S, Zon LI. Transplantation and in vivo imaging of multilineage engraftment in zebrafish bloodless mutants. Nat Immunol. 2003;4(12):1238–46. Epub 20031109. doi: 10.1038/ni1007. PubMed PMID: 14608381.

104. Harrold I, Carbonneau S, Moore BM, Nguyen G, Anderson NM, Saini AS, Kanki JP, Jette CA, Feng H. Efficient transgenesis mediated by pigmentation rescue in zebrafish. Biotechniques. 2016;60(1):13–20. Epub 20160101. doi: 10.2144/000114368. PubMed PMID: 26757807; PMCID: PMC4768720.

