## Supplementary material for "RNA Binding Protein Khdrbs1 Regulates Hematopoietic Stem and Progenitor Cell Emergence via Splicing": Karp_SuppFig_Leg

### SUPPLEMENTAL FIGURES AND LEGENDS

**Figure list:**

Figure S1 (Supplemental figure accompanying Fig 4): Khdrbs1 regulates other HE formation and differentiation into T-cells

Figure S2 (Supplemental figure accompanying Fig 4): Khdrbs1 regulates the myeloid and erythroid lineages

Figure S3 (Supplemental figure accompanying Fig 5): Other splicing events contain enriched Khdrbs1 binding sites

Figure S4 (Supplemental figure accompanying Fig 5): Splice site strength does not correlate with the presence of Khdrbs1 binding site

Figure S5 (Supplemental figure accompanying Fig 6): Khdrbs1 regulates HSPC formation via a splicing mechanism

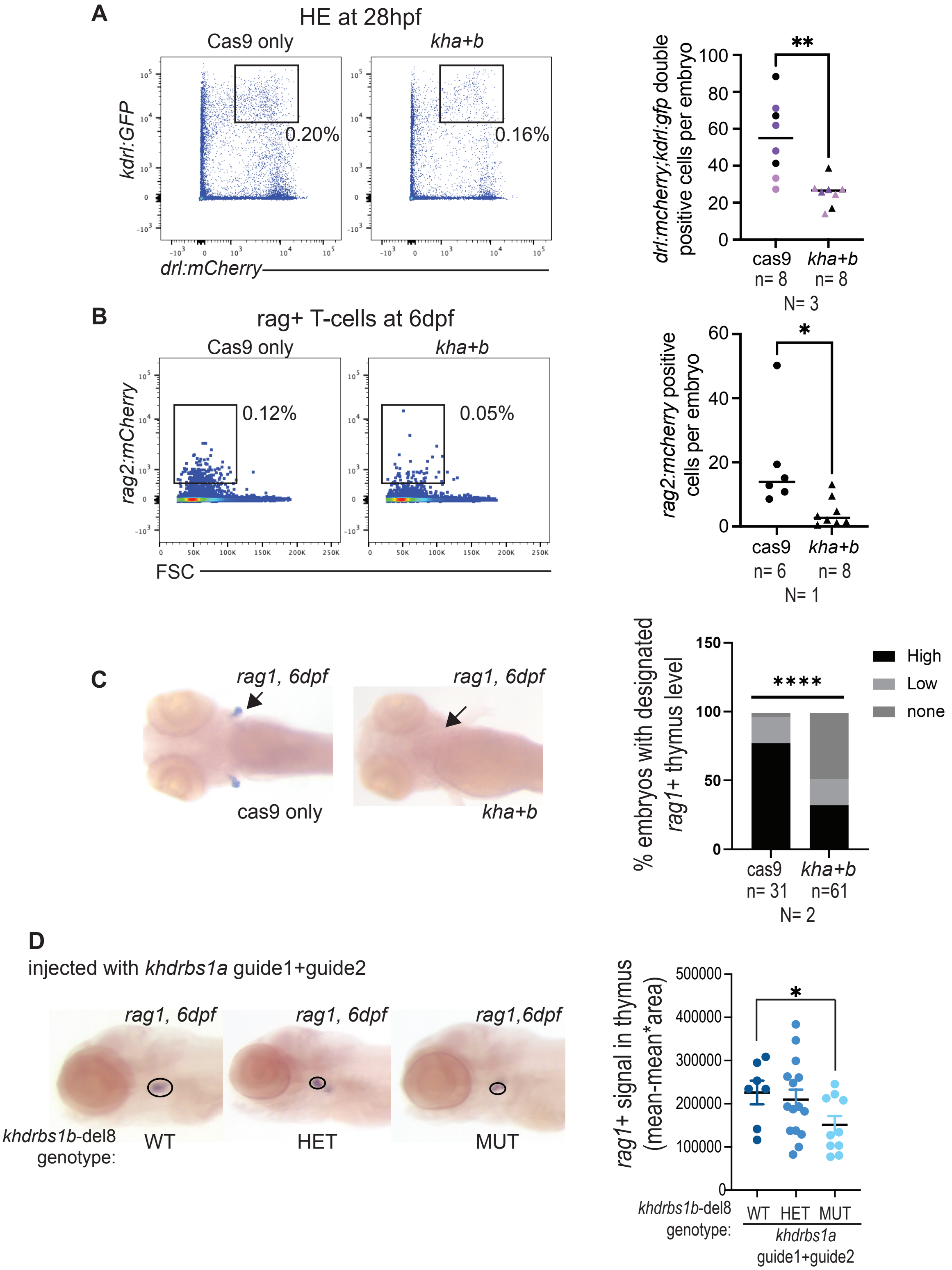


**Figure S1: Khdrbs1 regulates other HE formation and differentiation into T-cells.**

[**A**] Flow cytometry analysis using the LSRII instrument of *kdrl:*GFP and *drl*:mCherry expression in *khdrbs1a+b* crispants (*kha+b*) and cas9 only control at 28dpf. Representative flow plots are shown on the left and quantification is shown on the right. Each dot represents a pool of 4-10 embryos analyzed together and then graphed for numbers of *drl:*mCherry*;kdrl:*GFP double positive cells per embryo. Analysis done using FlowJo. [**B**] Flow cytometry analysis using the LSRII instrument of *rag2:*cherry expression in *khdrbs1a+b* crispants and cas9 only control at 6dpf. Representative flow plots are shown on the left and quantification is shown on the right. Each dot represents a pool of 10 embryos analyzed together and then graphed for numbers of *rag2+* cells per embryo. Analysis done using FlowJo. **pvalue<0.01 calculated by a student t-test with welch's correction. Different colors represent independent replicates. [**C**] T-cell levels in *khdrbs1a+b* crispants and cas9 only control were assessed by *rag1 in situ* hybridization at 6dpf. Representative images are shown on the left and quantification of *rag1* expression within the thymus shown to the right. ****pvalue<0.0001 calculated by a student t-test with welch's correction. Images were binned based on high expression of *rag1* (cas9 shown as an example), low expression of *rag1*, and no expression of *rag1* (*khdrbs1a+b* crispants shown as an example). [**D**] T-cell levels in *khdrbs1b-del8* wildtype (WT), heterozygous (HET), and homozygous (MUT) mutants injected with two sgRNA for *khdrbs1a* crispants were assessed by *rag1 in situ* hybridization at 6dpf. Representative images are shown on the left and quantification of *rag1* expression within the thymus shown to the right. *pvalue<0.05 calculated by a student t-test with welch's correction.

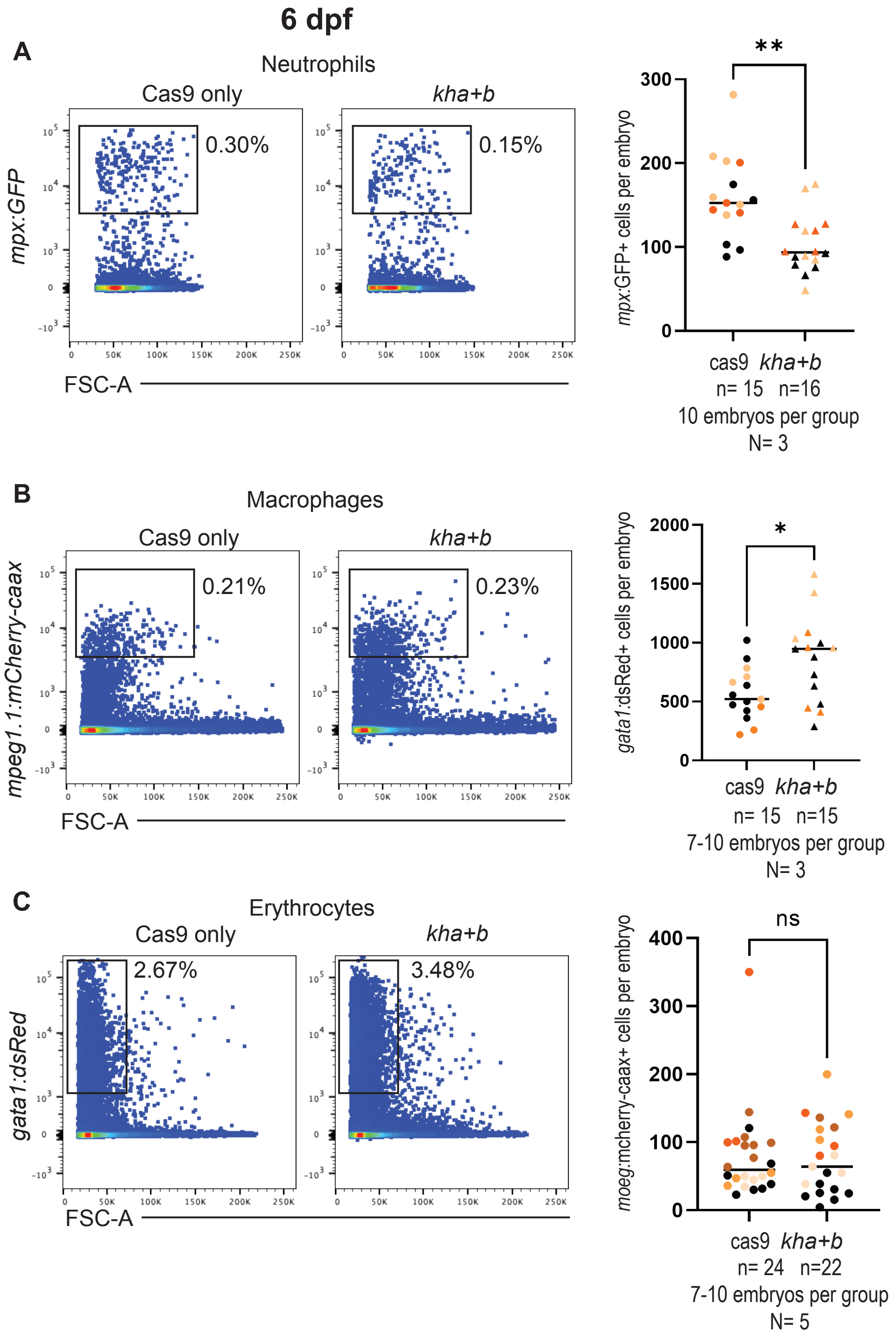


**Figure S2: Khdrbs1 regulates the myeloid and erythroid lineages.**

[**A-C**] Flow cytometry analysis using the LSRII instrument of *mpx:*GFP expression in *khdrbs1a+b* crispants (*kha+b*) and cas9 only control at 6dpf. Each dot represents a pool of 7-10 embryos analyzed together and then graphed for numbers of *mpx:*GFP+ (**A**), *mpeg:*mcherry-caax+ (**B**), and *gata1:*dsRed+ (**C**) cells per embryo. Analysis done using FlowJo. **pvalue<0.01 calculated by t-test with welch's correction. n.s indicates non significance. Different colors represent independent replicates.

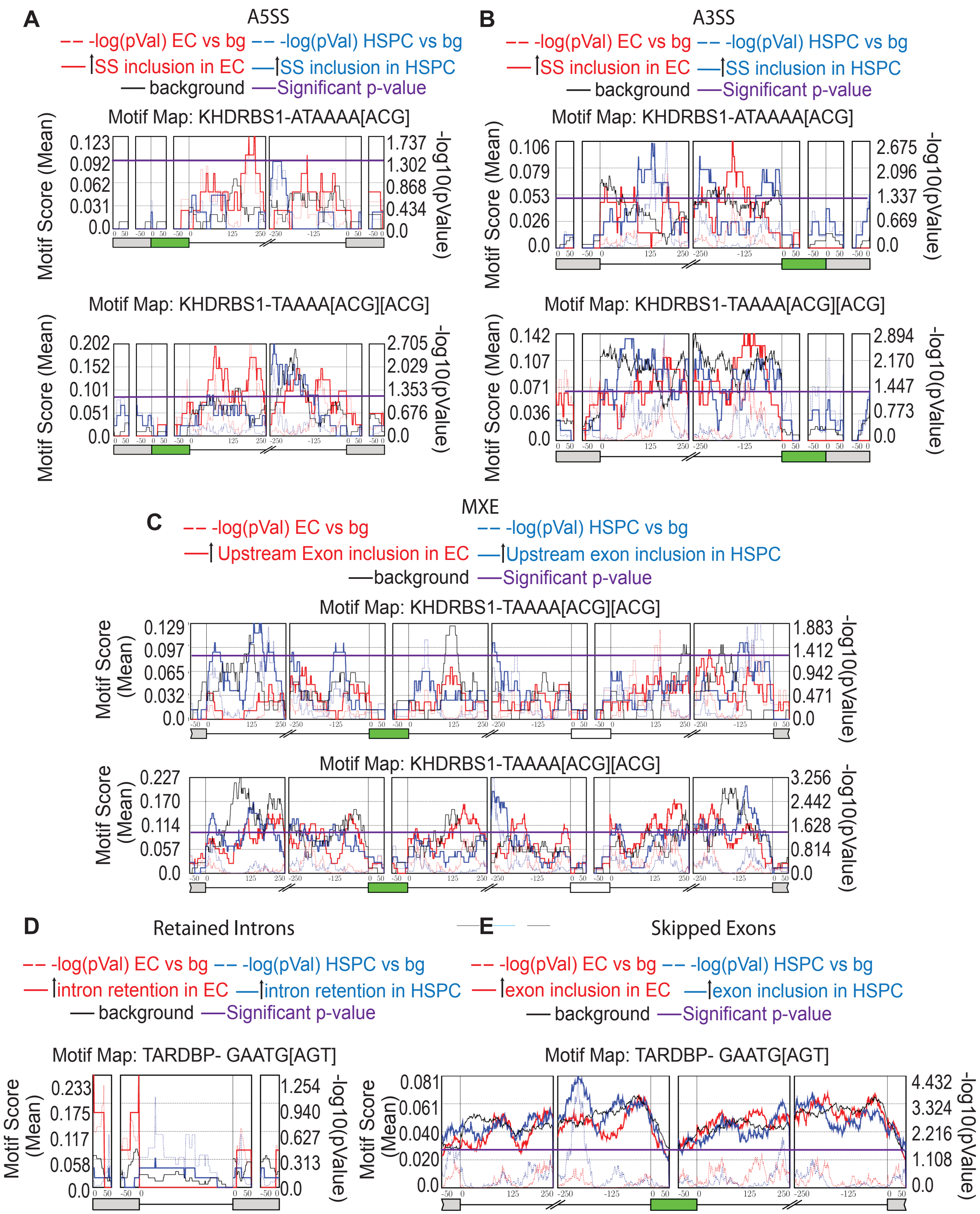


**Figure S3: Other splicing events contain enriched Khdrbs1 binding sites.**

[**A-C**] Metaplot of Khdrbs1 motif analysis of A5SS [**A**], A3SS [**B**], and MXE [**C**] from rMAPS[55] using significantly differentially spliced events in data from Xue et al., 2019[47] from Figure 1. Location of two different Khdrbs1 binding motifs were analyzed. Significance cutoff for differentially spiced events is FDR<0.05 and ΔPSI>10% Solid line indicates motif score and dotted line indicated negative log(p-value). Background events were taken from non-significant values of FDR=1. Significance was defined as p-value <0.05 (purple line). [**D-E**] Metaplot of Tardbp motif analysis of RI [D] and SE [E], from rMAPS[55] using significantly differentially spliced events in data from Xue et al., 2019[47]. Background events were taken from non-significant values of FDR =1. Significance was defined as p-value <0.05 (purple line).

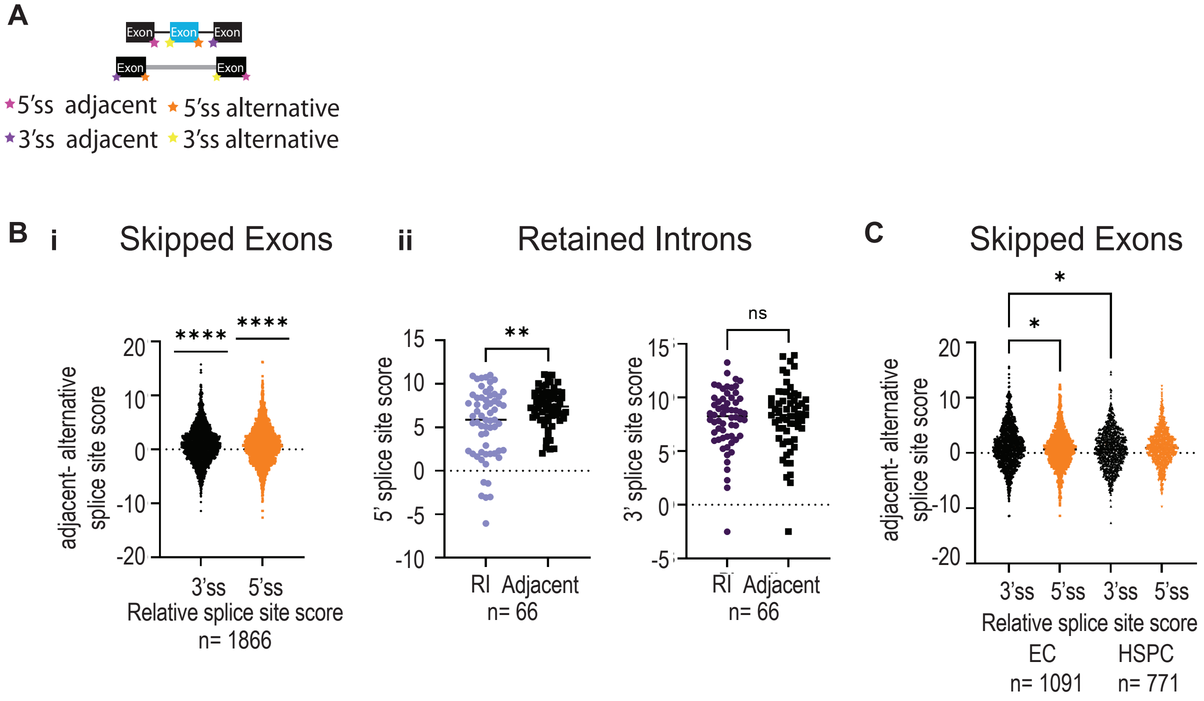


**Figure S4: Splice site strength does not correlate with the presence of Khdrbs1 binding site.**

[**A**] Schematic representing 5'ss and 3'ss used for alternative and adjacent exons for SE and retained introns and adjacent exons for RI. [**B**] Splice site strength analysis using MaxEntScan[99] for SE (i) and RI (ii) for all significantly differentially spliced events in Xue et al. (FDR<0.05 and ΔPSI>10%). Statistical test used was a one-sided t-test; **** represents p-value <0.0001.

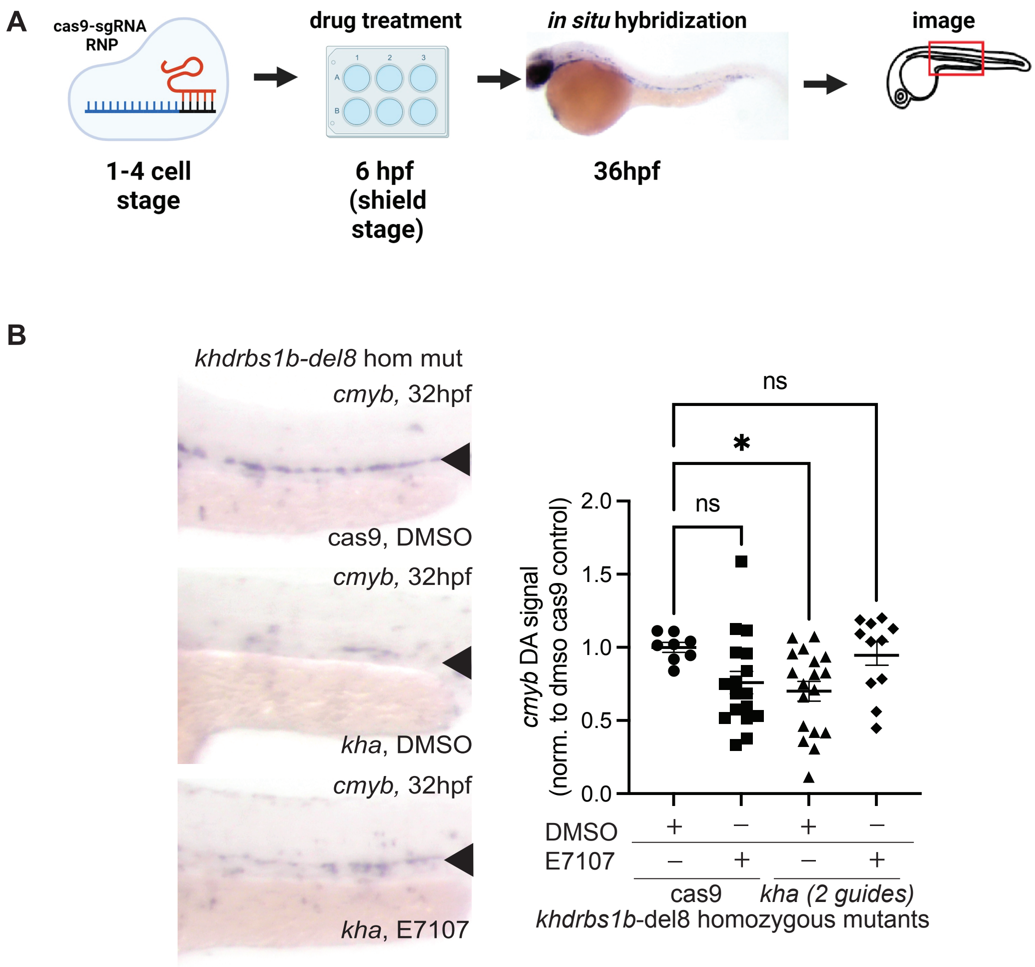


**Figure S5: Khdrbs1 regulates HSPC formation via a splicing mechanism.**

[**A**] Schematic representation of workflow studying the effect of the splicing modulator E7107 on HSPC formation in *khdrbs1b* homozygous mutants mutagenized for *khdrbs1a*. Two guide RNAs for *khdrbs1a* or cas9 as a control were injected into an intercross of *khdrbs1b-del8* heterozygous animals. Embryos were treated from 6-36hpf with either 15µM E7107 or DMSO. *In situ* hybridization were performed at 34hpf using a *cmyb* probe and then imaged with dissecting scope with 8x magnification. [**B**] Quantification and images of cas9 only and *khdrbs1a* crispant (*kha*) *khdrbs1b-del8* homozygous mutant embryos at 34hpf with *in situ* hybridization of *cmyb* probe. Images were analyzed by image j after conversion to 8 bit and inversion of colors. Analysis was performed using one-way ANOVA. *p-value <0.05 and n.s represents non-significant p-value of >0.05.
