## Supplementary material for "RNA Binding Protein Khdrbs1 Regulates Hematopoietic Stem and Progenitor Cell Emergence via Splicing": Methods

Address: 1300 Morris Park Avenue, Bronx, NY 10461

### STAR METHODS

### RESOURCE AVAILABILITY

#### *Lead contact*

#### *Material availability*

Animal models generated in this paper will be shared freely upon request to the Lead Contact.

***Data availability***

- Microscopy data reported in this paper will be shared by the lead contact upon request.
- This paper does not report original code.
- Any additional information required to reanalyze the data reported in this work is available from the Lead Contact upon request.

### EXPERIMENTAL MODEL AND SUBJECT DETAILS

#### Zebrafish Maintenance and Handling

Zebrafish were maintained as described[75]. All fish were maintained by protocols approved according to institutional animal care and use committee in accordance with Albert Einstein College of Medicine research guidelines.

#### Zebrafish Lines

The strains of animals used include wild type (AB), casper, Tg(*kdrl*:GFP)[54], Tg(*mpx*:gfp)[76], Tg(*mpeg1.1*:mcherry-caax[77]), Tg(*rag2*:mcherry)[78], and *esrp1* and *esrp2* double mutants[79] maintained as double heterozygous adults. A new *khdrbs1b* mutant containing an 8bp deletion was generated using CRISPR-Cas9 mutagenesis with the sgRNA used in the RBP screen. Embryos were collected and staged as previously described[80].

#### Whole-mount in situ hybridization.

*In situ* hybridization (ISH) for *notch1b*[81], *runx1*[82], *cmyb*[83], and *rag1*[84] was performed, with minor modifications, as described previously[85]. Embryos (28 or 36-40hpf) and larvae (6dpf) were bleached with 0.8% KOH, 0.9% H2O2, and 0.1% Tween 20 for 5-7.5 min or 25 minutes, respectively. They were then permeabilized in Proteinase K (Roche) for 5-7.5 minutes(embryos) or 20 minutes (larvae). After ISH, embryos and larvae were stored in 4% paraformaldehyde (PFA) until imaged.

**Microscopy and Image Analysis**

ISH embryos were placed in 100% glycerol and imaged using Zeiss Stereo Discovery.V8 microscope at 8x magnification. ISH signal was quantified as previously described [Dobrzycki et al. PMID: 29535102]. Briefly, imageJ[86] was used to invert the signal, convert to 8-bit, and measure pixel intensity in the area of interest and in the background. For ISH images displayed in the manuscript, white balance adjustments were performed in Adobe Photoshop 2024.

For live imaging, Tg(*rag2:*mcherry) larvae were embedded in 3% methylcellulose dissolved in E3 embryo water (NaCl, KCl, $MgSO_4 \cdot 7H_2O$, $CaCl_2 \cdot 2H_2O$), then they were imaged using a Zeiss Axiovert inverted microscope at 5x magnification (6dpf larvae) and 10x magnification (4dpf and 5dpf larvae). Fluorescence of the mCherry signal within the thymus was quantified using imageJ.[86] For display purposes, images were adjusted for display in Adobe Photoshop 2024 using the brightness/contrast tool by 100 twice where the adjustment scale ranges -100 to 100.

**Sample Collection and Dataset Utilization**

Published RNA sequencing datasets were utilized for differential splicing and gene expression analyses. For the HE vs EC comparison, we utilized Lefkopoulos et al.[45]. In that study, hemogenic endothelium (HE) and endothelial cells (EC) were isolated at 26 hours post-fertilization (hpf). HE were defined as cells expressing both *cmyb:GFP* and *kdrl:CFP* expression and EC were defined as positive for *kdrl:GFP* or *kdrl:CFP*. For the HE/HSPC vs EC comparison, we utilized Li et al In that study, cells were isolated from 36 hpf embryos with HE/hematopoietic stem/progenitor cells (HSPCs) defined as cells expressing *runx1en:gfp* and *kdrl:mcherry* and EC defined as cells expressing *kdrl:mcherry*. For the HSPC vs EC comparison, we utilized Xue et al[47]. In that study, cells were isolated from 36 hpf embryos with HSPC defined as cells expressing *cd41:gfp* and EC defined as cells expressing *kdrl:mcherry*. Fastq files were downloaded and were mapped to the zebrafish genome version GRCz10/danRer10 with STAR[87] aligner (version −2.7.10b). Alignment was performed using the settings: --outSAMtype BAM SortedByCoordinate --outSAMstrandField intronMotif --outSAMattributes All --outFilterMultimapNmax 1 --outFilterScoreMinOverLread 0.51 --outFilterMatchNminOverLread 0.51 --

outFilterMismatchNmax 6 --alignIntronMax 50000 --sjdbGTFfile $gtf --quantMode TranscriptomeSAM GeneCounts --twopassMode Basic

**Differential Splicing Analysis**

RNA sequencing data were analyzed using rMATS[48] version 4.1.0 to detect differential splicing events. rMATS settings utilized were -t paired --variable-read-length. Read length was 150bp for Xue et al.[47] and 75bp for Li et al. and Lefkopoulos et al.[45]. Significant splicing events were defined by false discovery rate (FDR) < 0.05 and a change in Percent Spliced In (ΔPSI) > 10%. Types of splicing events analyzed included skipped exon (SE), retained intron (RI), alternative 5' splice site (A5SS), alternative 3' splice site (A3SS), and mutually exclusive exons (MXE).

**Differential Gene Expression Analysis**

Gene counts for each sample were generated using featureCounts[88] version 2.0.1 with settings -C -B -p. Differential gene expression analysis was conducted using the DESeq2[89] package 1.42.0 using R version 4.3.2. Genes with a log2FoldChange > 0.32 and an adjusted p-value < 0.05 were considered significantly differentially expressed between EC and HSPC or HE samples.

To determine which of the differentially expressed genes are RBPs, gene symbols were converted from zebrafish to human using BioMart [[90, 91]] and then human gene names were compared to existing RBP databases. Databases utilized were RBP2GO[49] and RBPDB[74]. Of the 67 differentially expressed RBP, those with splicing factor functions were then identified using a literature search.

**Pathway Analysis**

Pathway enrichment analysis of differentially expressed genes (DEGs) was performed using Metascape[92] with standard metascape settings. Pathways were analyzed for each set of differentially expressed or differentially spliced genes for all genes that were higher in HSPC or HE, or higher in EC. Metascape groups pathways into one summary pathway based on similarities. For each summary pathway, one pathway was selected for final presentation.

**Functional Screening of RBPs**

RBPs were grouped based on differential gene expression levels and functional annotations. For genes with higher inclusion in ECs, Group 1 included all KH domain containing genes except for *nova1: qkia, khsrp,* and *khdrbs1a.* The paralog of *khdrbs1a, khdrbs1b,* was included as mutation of one paralog often results in compensation by the other paralog. Group 2 included all HNRNP genes: *hnrnpaba, hnrnpub, hnrnpua,* and *hnrnpa0a.* Group 3 contained those that were not included in the other two groups: *nova1, mbnl1, tardbp, rbm15b.* All genes with higher inclusion in HSPCs (*rnpc3, mbnl1,* and *zranb2)* were tested individually. For *esrp1* and *esrp2,* mutant zebrafish lines were acquired from the Liao lab[79] and were maintained as double heterozygous animals.

For all crispants, two gRNAs were designed and tested for each RBP gene using a T7 endonuclease assay to assess mutagenesis efficiency[93]S Functional screening for hematopoietic stem and progenitor cell (HSPC) formation was conducted via *in situ* hybridization for *cmyb* expression at 36 hpf. Quantification of expression levels was performed using ImageJ as described above.

**Injecting CRISPR gRNAs**

To generate fully functional CRISPR RNA for inducing Cas9-based mutagenesis, gRNAs were combined with tracrRNA (IDT), nuclease-free duplex buffer (IDT), and Cas9 protein (IDT). Assembled RNP complexes were injected into zebrafish embryos at the 1-4 cell stage. Mutagenesis efficiency was assessed using a T7 endonuclease assay[94]. For each gene, the best gRNA was utilized for the pooled screened. For injections, 0.5 nL droplets of gRNA::tracrRNA::Cas9 RNP complexes were injected. When four gRNA::tracrRNA were injected simultaneously, the concentration of each gRNA::tracrRNA in the injection solution is ~125ng/ml and ~187.5ng/ml for two crRNA::tracrRNA.

**Designing CRISPR gRNAs**

CRISPR gRNAs were designed using CHOPCHOP[95-97]. Target sites were selected based on presence of PAM sequences and minimal off-target potential. The different guides were selected in different exons. Guides tested are listed in Table S5.

**T7 endonuclease(T7E) assay:**

For each gRNA, the sequences flanking the targeted region were amplified by PCR. Following PCR (35 cycles at 58°C), DNA strands in the samples were separated by incubating in a thermocycler at 98°C for 3 min and reannealed by cooling slowly in an unplugged machine for 2 hrs. Samples were incubated in T7E (NEB) and 1XNEB Buffer 2 at 37°C for 15 min. Digested and undigested PCR products were run on a 2-3% agarose gel. Mutagenesis was confirmed based on the presence of T7E cut bands in the crispant group relative to Cas9 only controls. Primers used for PCR are listed in Table S5.

***esrp1/esrp2* testing**

Genotyping for *esrp1$^{fb401}$* WT and mutant alleles was performed with a custom Taqman single nucleotide polymorphism (SNP) probe and TaqMan universal PCR mastermix (Thermo Fischer) using a StepOne Plus Real Time PCR machine. The *esrp1-/-* embryos were then genotyped for *esrp2$^{fb402}$* by T7E assay using *esrp2* specific primers. In T7E-negative embryos, *esrp2* WT DNA was spiked in to form hybrid substrates with mutant DNA, which will distinguish WT from mutant embryos. Primers for amplifying the *esrp2* mutant flanking region are listed in Table S5.

**Drug treatment**

Zebrafish embryos were treated with 30 mM of the splicing modulator E7107 [gift from H3 Biomedicine] or DMSO. Embryos were treated from 6-36 hpf. The drug treatments were washed out and the embryos were dechorionated with pronase (Roche) followed by fixation with 4% PFA.

**Motif Analysis**

Motif analysis for splicing factors was performed using RNA map analysis and plotting server for alternative exon regulation (rMAPS)[55]. Significantly differentially spliced genes (FDR<0.05 ΔPSI>=10%) from data from Xue et al.[47], were analyzed as higher in EC or higher in HSPC against background events. Background events were those identified by rMATS within the same dataset as FDR = 1. SE, RI, MXE, A5SS, and A3SS were all analyzed.

Individual splicing events were analyzed for the presence of a Khdrbs1 splicing motif (ATAAAA[ACG]) using find individual motif occurrences (FIMO)[98] with a p-value <0.001. For

RI, the input sequences were the retained introns and flanking exons, while for SE the sequences were the alternative exons and 250 bp in the upstream and downstream introns.

**Splice Site Strength**

Splice site strength analysis was conducted using the maximum entropy model of MaxEntScan[99] for SE and RI events. For RI, the RI 5'ss was compared to the downstream 5'ss and the RI 3'ss was compared to the upstream 3'ss. For SE, the alternative exon 5'ss was compared to the upstream 5'ss and the alternative exon 3'ss was compared to the downstream 3'ss. Since alternative exons are in direct competition with adjacent exons, for SE, relative splice site scores were calculated by subtracting the alternative exon splice site score from adjacent.

**Flow Cytometry and Staining:**

Embryos were removed from their chorions using pronase (Roche), pooled, and manually dissociated using a sterile razor blade. To further homogenize the samples, they were incubated with 1× Dulbecco phosphate-buffered saline (PBS; D-PBS) (Life Technologies) supplemented with a 1/65 dilution of 5 mg/mL Liberase (Roche) and incubated at 37°C for 6 min for 28-40 hpf embryos and 10 min for 6dpf larvae. Liberase was neutralized using 5% fetal bovine serum (FBS)(Life Technologies). The cells were filtered through a 40-mm cell strainer (Falcon)were resuspended in FACS Buffer (0.9× D-PBS, 5% FBS, 1% Penn/Strep [Life Technologies]) containing 1mg/mL 4′,6-Diamidino-2-phenylindole (DAPI) to identify dead cells. Flow cytometry analysis was performed using an LSRII Flow cytometer (BD BioSciences) and FlowJo version 10.0.08.

**QUANTIFICATION AND STATISTICAL ANALYSIS**

Two tailed Student T-test with Welch's correction was used for pairwise comparison. Analysis of variance (ANOVA) was used when more than two samples were analyzed. For directional testing, one-sided T-test was used. Statistical analyses were performed using GraphPad Prism. Error bars indicate standard deviation from mean, or as specified. ns, not significant; * $p<0.05$; ** $p<0.01$; *** $p<0.001$; **** $p<0.0001$.
