## Supplementary material for "RNA Binding Protein Khdrbs1 Regulates Hematopoietic Stem and Progenitor Cell Emergence via Splicing": KeyResources

Address: 1300 Morris Park Avenue, Bronx, NY 10461

**KEY RESOURCES TABLE**

| REAGENT or RESOURCE | SOURCE | IDENTIFIER |
|---|---|---|
| Antibodies | | |
| Anti-digoxigenin antibody | Roche | 11093274910 |
| Bacterial and virus strains | | |
| Biological samples | | |
| Chemicals, peptides, and recombinant proteins | | |

| | | |
|---|---|---|
| Proteinase K | Roche | RPROTK-RO |
| Paraformaldehyde | Affymetrix | J19943.K2 |
| Glycerol | Sigma | G5516 |
| Cas9 | IDT | 1081058 |
| Pronase | Roche | PRON-RO |
| PBS;D-PBS | Life Technologies | D8662 |
| Tween20 | Sigma | P1379 |
| Liberase | Roche | LIBTM-RO |
| Penicillin-Streptomycin | Life Technologies | 15140122 |
| Methanol | Fisher | BPA452SK4 |
| 4’6-Diamidino-2-phenylindole | Sigma | D8417 |
| Digoxigenin-NTPs | Roche | DIGUTP-RO |
| Bovine Serum Albumin | Sigma | A7037 |
| T7 RNA polymerase | Roche | RPOLT7-RO |
| RNase Inhibitor I | Roche | RNAINH-RO |
| Dimethyl sulfoxide | Sigma | D5879 |
| E7107 | Gift from H3 Biomedicine | NA |
| DNase I | Roche | 04716728001 |
| Critical commercial assays | | |
| Purelink PCR purification Kit | Thermo Fisher | K310002 |

| | | |
|---|---|---|
| Deposited data | | |
| HEvsEC | Lefkopoulos et al., 2020[45] | SRA: PRJNA578896 (BioProject ID) |
| HE/HSPCvsEC | Li et al., 2022[46] | GSE189072. |
| HSPCvsEC | Xue et al., 2019[100] | GSE120581 |
| Experimental models: Cell lines | | |
| Experimental models: Organisms/strains | | |
| Wild Type AB | | |
| Tg(*kdrl*:GFP) | Jin et al., 2005[54] | ZFIN ID: ZDB-TGCONSTRCT-070529-1 |
| Tg(*mpx*:gfp) | Renshaw et al., 2006[101] | ZDB-TGCONSTRCT-070118–1 |
| Tg(*mpeg1.1*:mcherry-caax) | Ellet et al., 2011[102] | ZFIN ID: ZDB-PUB-101122-22 |
| Tg(*drl*:mcherry) | Sanchez-Iranzo et al., 2018 | ZDB-TGCONSTRCT-171031–8 |
| Tg(*gata1:*dsRED) | Traver et al., 2003[103] | ZDB-TGCONSTRCT-070117-38 |

| | | |
|---|---|---|
| Tg(*rag2*:mcherry) | Harrold et al., 2016[104] | ZDB-TGCONSTRCT-160329-2 |
| *khdrbs1b*$^{-/-}$ | This paper | NA |
| *esrp1 esrp2*[79] | Carroll et al., 2006[79] | ZDB-ALT-211104-3 ZDB-ALT-211104-4 |
| Oligonucleotides | | |
| tracrRNA | IDT | 1072532 |
| Custom crRNA | IDT | See table 5 |
| Recombinant DNA | | |
| Software and algorithms | | |
| STAR 2.7.10b[87] | Dobin et al., 2013 | **https://github.com/alexdobin/STAR/releases** |
| rMATS[48] | Shen et al., 2014 | https://github.com/Xinglab/rmats-turbo/releases/tag/v4.1.0 |
| featureCounts 2.0.1[88] | Liao et al., 2014 | http://subread.sourceforge.net |
| DESeq2 1.42.0[89] | Anders et al., 2010 | https://bioconductor.org/packages/release/bioc/html/DESeq2.html |

| | | |
|---|---|---|
| Metascape[92] | Zhou et al., 2019 | https://metascape.org/ |
| rMAPS[55] | Park et al., 2016 | http://rmaps.cecsresearch.org/ |
| FIMO[98] | Grant et al., 2011 | https://meme-suite.org/meme/doc/fimo.html |
| MaxEntScan[99] | Yeo et al., 2004 | http://hollywood.mit.edu/burgelab/maxent/Xmaxentscan_scoreseq.html |
| Graphpad Prism 9.4.1 | Graphpad | https://www.graphpad.com/ |
| CHOPCHOP[96] | Montague et al., 2014 | **https://chopchop.rc.fas.harvard.edu** |
| BioMart[91] | Smedley et al., 2009 | http://www.biomart.org |
